# Spontaneously regenerative corticospinal neurons in mice

**DOI:** 10.1101/2024.09.09.612115

**Authors:** Benjamin W Fait, Bianca Cotto, Tatsuya C. Murakami, Michael Hagemann-Jensen, Huiqing Zhan, Corinne Freivald, Isadora Turbek, Yuan Gao, Zizhen Yao, Sharon W. Way, Hongkui Zeng, Bosiljka Tasic, Oswald Steward, Nathaniel Heintz, Eric F Schmidt

## Abstract

The spinal cord receives inputs from the cortex via corticospinal neurons (CSNs). While predominantly a contralateral projection, a less-investigated minority of its axons terminate in the ipsilateral spinal cord. We analyzed the spatial and molecular properties of these ipsilateral axons and their post-synaptic targets in mice and found they project primarily to the ventral horn, including directly to motor neurons. Barcode-based reconstruction of the ipsilateral axons revealed a class of primarily bilaterally-projecting CSNs with a distinct cortical distribution. The molecular properties of these ipsilaterally-projecting CSNs (IP-CSNs) are strikingly similar to the previously described molecular signature of embryonic-like regenerating CSNs. Finally, we show that IP-CSNs are spontaneously regenerative after spinal cord injury. The discovery of a class of spontaneously regenerative CSNs may prove valuable to the study of spinal cord injury. Additionally, this work suggests that the retention of juvenile-like characteristics may be a widespread phenomenon in adult nervous systems.

## Introduction

The mammalian spinal cord receives input from the motor cortex via the corticospinal tract (CST).^1–3^ The CST comprises corticospinal neuron (CSN) axons that begin in Layer 5b of motor cortex and can extend up to one meter to reach targets in the caudal spinal cord in humans.^4^ CSNs regulate the spinal cord’s sensorimotor function; however, the extent of their impact on motor control differs between species.^5,6^ In macaques, lesion of the pyramidal tract produces immediate limb paralysis.^7,8^ In rodents, their genetic ablation profoundly impacts skilled-reaching ability but does not produce paralysis.^9^

In almost all examined mammals,^10–12^ corticospinal axons primarily terminate contralateral to their cell bodies of origin.^1,13–15^ Understanding the contralateral organization of the descending tracts is one of the oldest topics in neuroscience.^16–18^ Writing of the tendency of forebrain injuries to produce neurological signs on the side opposite the injury, Aretaeus of Cappadocia (c. 100 CE) reasoned, “The cause of this is the interchange in the origin of nerves, for they do not pass along on the same side…but each of them passes over to the other side from that of its origin, decussating each other in the form of the letter *Χ*.”^19^

Despite the predominance of contralateral CSN termination, a minority of CSN axons terminate ipsilaterally. Studies using biotinylated dextran amine (BDA) to trace anterogradely from the primary motor cortex (M1) to the cervical spinal cord in macaques found that ∼2%−18% of total axons terminated ipsilaterally.^20,21^ Similar experiments labeling from M1 to the cervical spinal cord in mice detected ∼0.4−2.3% of synaptic boutons and ∼3.5−7% of total axons on the ipsilateral side of the spinal cord.^22–25^ In mice, disruption of several genes encoding axonal guidance molecules increased the number of ipsilateral projections.^24,26–31^ Ipsilateral axons are more abundant during development before activity-dependent pruning, and pharmacological inactivation of one hemisphere of the motor cortex during this period increases their abundance from the active hemisphere.^3,32^ There has been considerable examination of ipsilateral axons in unilateral cortical injury because of evidence that they are a potential circuit for the uninjured cortex to compensate for the damaged one.^33–35^ However, little is known about their anatomy and function in healthy animals.

Though sparse, these ipsilateral axons are a remarkable deviation from the norm. Brains have evolved only in bilaterally symmetric animals, so the midline is a foundational architectural principle of all central nervous systems.^36–38^ Accordingly, an intricate and redundant ensemble of molecular signals exerts tight control over midline crossing.^39^ The ipsilateral axons represent a consistent anomaly across many species at an evolutionarily ancient midline boundary that is under strict developmental control.

To better understand these ipsilateral axons, we first sought to characterize their anatomy with a diverse array of emerging methods in mouse molecular neuroscience. Using anterograde tracing methods, tissue clearing, and Smartseq3 single-nucleus RNA-sequencing (snRNA-seq), we found that ipsilateral CSN axons project to regions of the ventral horn, including directly to motor neurons. Barcode-based Multiplexed Analysis of Projections by Sequencing (MAPseq) of the CST revealed that the neurons contributing these axons primarily comprise a class of bilaterally projecting CSNs and represent a more substantial population of total CSNs than their sparse ipsilateral axons suggest. With retrograde tracing and tissue clearing, we found that the cell bodies of this neuronal population reside in distinct cortical regions, almost entirely absent from the caudo-lateral cortex. We deeply profiled their molecular characteristics using the viral implementation of Translating Ribosome Affinity Purification (vTRAP) and discovered a striking similarity to the embryonic-like molecular signature of regenerating corticospinal neurons. We noticed that the anatomical characteristics of IP-CSNs we had documented were shared with regenerating CSNs: in addition to molecular similarity, both share bilaterality, projection to ventral spinal cord regions, and motor neuronal connectivity. Given these similarities, we hypothesized that IP-CSNs might themselves have regenerative properties. Finally, we show that IP-CSNs are spontaneously regenerative.

## Results

### The ipsilateral axons of the CST project to ventral spinal cord regions

Examining the comparative spatial projections of the CST in various mammals has proven a valuable starting point for considering its function.^40,41^ Therefore, we sought to compare the post-synaptic targets of ipsilaterally and contralaterally terminating CSN axons in mice. To approach this question, we used AAV1-Cre anterograde monosynaptic tracing in Ai14^42^ (Cre-dependent tdTomato [tdT] reporter mice). This approach results in anterograde trans-synaptic transfer of the virus and Cre-mediated tdT fluorescence in the post-synaptic neurons of CSNs.^43,44^ AAV1-Cre was injected into the caudal forelimb area (CFA) of the motor cortex of Ai14 mice and allowed to express for five weeks **(Fig 1A−B**). We then analyzed 8,183 post-synaptic neurons in whole cervical spinal cords using CUBIC tissue clearing and light-sheet microscopy and compared the relative distributions of labeled cells on either side of the spinal cord.^45^

**Fig 1:**
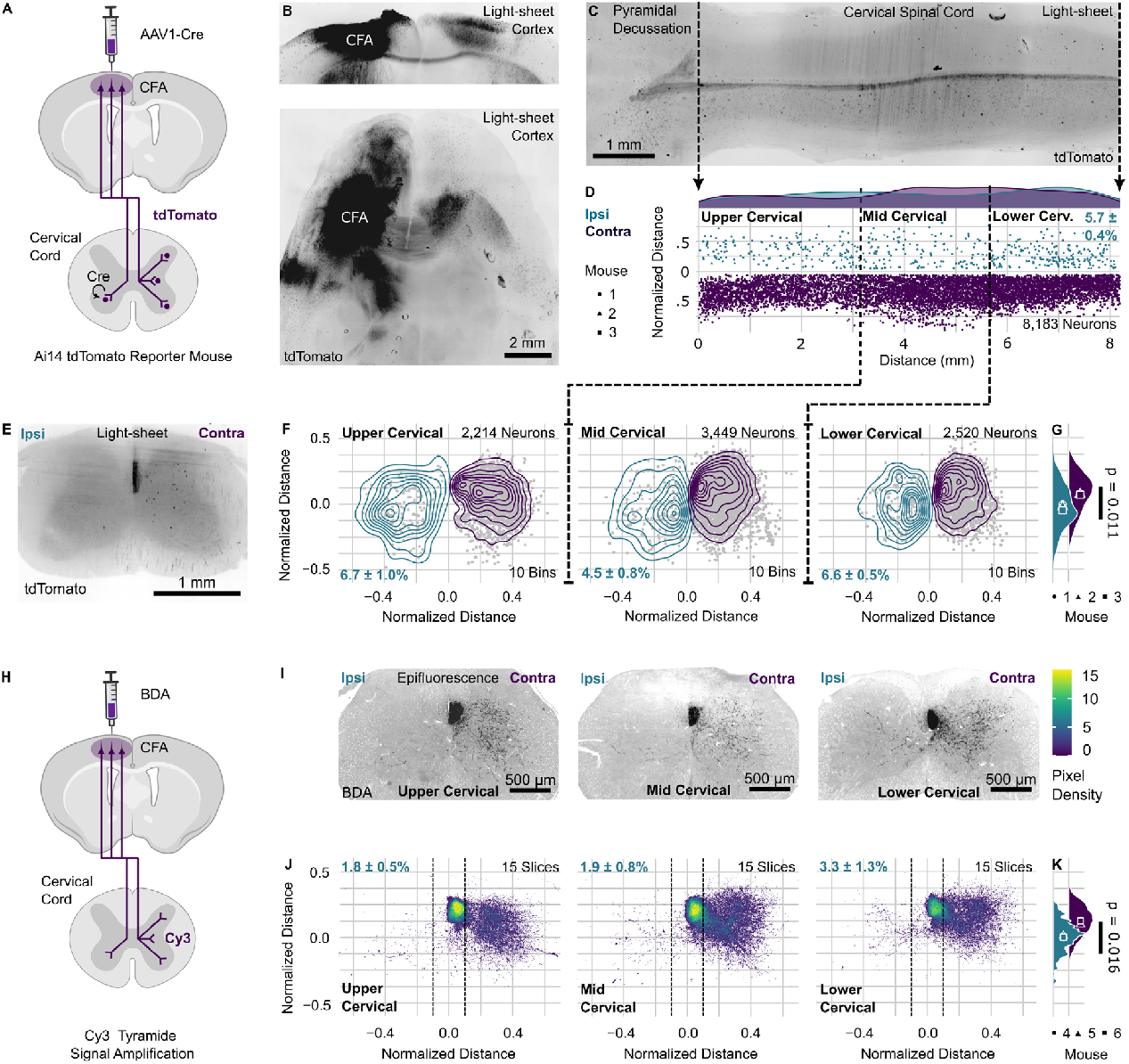
The ipsilaterally terminating axons of the CST project to ventral spinal cord regions. **(A)** A schematic of the AAV1-Cre monosynaptic anterograde tracing experiment (n = 3). **(B)** Maximum-intensity z-projections of a cleared injection site on the coronal and horizontal planes with native tdT fluorescence. **(C)** A maximum-intensity z-projection of the cleared cervical spinal cord on the coronal plane with native tdT fluorescence. The brightness of representative images was adjusted so that tissue was visible via autofluorescence. **(D)** A scatter plot of the 8,183 tdT(+) neurons on the coronal plane, with ipsilateral in teal and contralateral in purple. Above, density plots show the rostro-caudal distributions of neurons. **(E)** A 100 μm maximum-intensity z-projection of the cervical spinal cord on the horizontal plane with tdT(+) neurons. **(F)** Three scatter plots on the horizontal plane of annotated neurons in gray, proceeding left to right from high to low cervical regions as indicated by the dotted lines in (D). Coordinates are normalized to the height and width of the spinal cord grey matter. Density plots of their distributions (10 bins) are overlaid, with ipsilateral in teal and contralateral in purple. **(G)** Ridgeline plots show the ventral distribution of plotted neurons across the cervical spinal cord (p = 0.011, n = 3, paired two-sided t-test), with ipsilateral in teal and contralateral in purple. **(H)** A schematic of the BDA tracing experiment (n = 3). **(I)** Representative images of the BDA-labeled (Cy3) corticospinal axons on the horizontal plane with ipsilateral axons on the left side and contralateral on the right. **(J)** Three heatmaps on the horizontal plane show the distribution of BDA-labeled axons from upper to lower cervical regions. The analysis was conducted on regions lateral to the dotted lines. The data were normalized to the length and width of the spinal cord grey matter. **(K)** Ridgeline plots show the distribution of all annotated pixels, with ipsilateral in teal and contralateral in purple (p = 0.016, n = 3, paired two-sided t-test).

Throughout the cervical spinal cord, 5.7 ± 0.4% of postsynaptically labeled tdT(+) neurons were on the ipsilateral side **(Fig 1C−D)**. These were distributed similarly along the rostro-caudal axis as the contralateral neurons (p = 0.67, n = 3, paired two-sided t-test) as well as on the medial-lateral axis (p = 0.10, n = 3, paired two-sided t-test), but tdT(+) neurons on the ipsilateral side were located more ventrally than their contralateral counterparts (p = 0.011, n = 3, paired two-sided t-test) **(Fig 1E−G)**. We confirmed this finding using anterograde BDA tracing from the CFA to label CSN axons on either side of the spinal cord **(Fig 1H)**. Of all BDA-labeled pixels in the cervical spinal cord, 2.4 ± 0.9% were on the ipsilateral side. As previously reported,^46^ we observed some axons descending in the ipsilateral dorsal funiculus; however, we did not observe axons exiting it. Instead, ipsilaterally terminating axons appear to reach the ipsilateral side of the spinal cord primarily by “re-crossing” via the spinal cord commissure from the contralateral side **(Fig 1I)**. The distribution of BDA-labeled axons was similar to the distribution of post-synaptic neurons, with the ipsilateral axons located more ventrally (p = 0.016, n = 3, paired two-sided t-test) **(Fig 1I−K)**.

In the broadest terms, sensory function is associated with the dorsal laminae of the spinal cord, while motor function is associated with the ventral laminae.^47^ These data suggest that ipsilateral CSN axons project to neuronal populations involved in motor control.

### Comprehensive transcriptomic characterization of postsynaptic cells targeted by ipsilateral and contralateral CSN axons

To better understand the cellular identity of projection targets of the ipsilateral and contralateral corticospinal axons, we repeated our AAV1-Cre monosynaptic tracing approach using Isolation of Nuclei Tagged in Specific Cell Types (IN-TACT) reporter mice, in which Cre-mediated recombination results in nuclear sfGFP fluorescence in post-synaptic neurons.^48,49^ Labeled nuclei from each side of the spinal cord were individually sorted and their transcriptomes were analyzed with Smart-seq3 snRNA-Seq^50^ **(Fig 2A, Fig S1A−D)**. Consistent with our prior experiments, we found that 4.3 ± 0.8% of total sfGFP(+) nuclei were located in the ipsilateral spinal cord samples **(Fig S1B)**.

**Fig 2:**
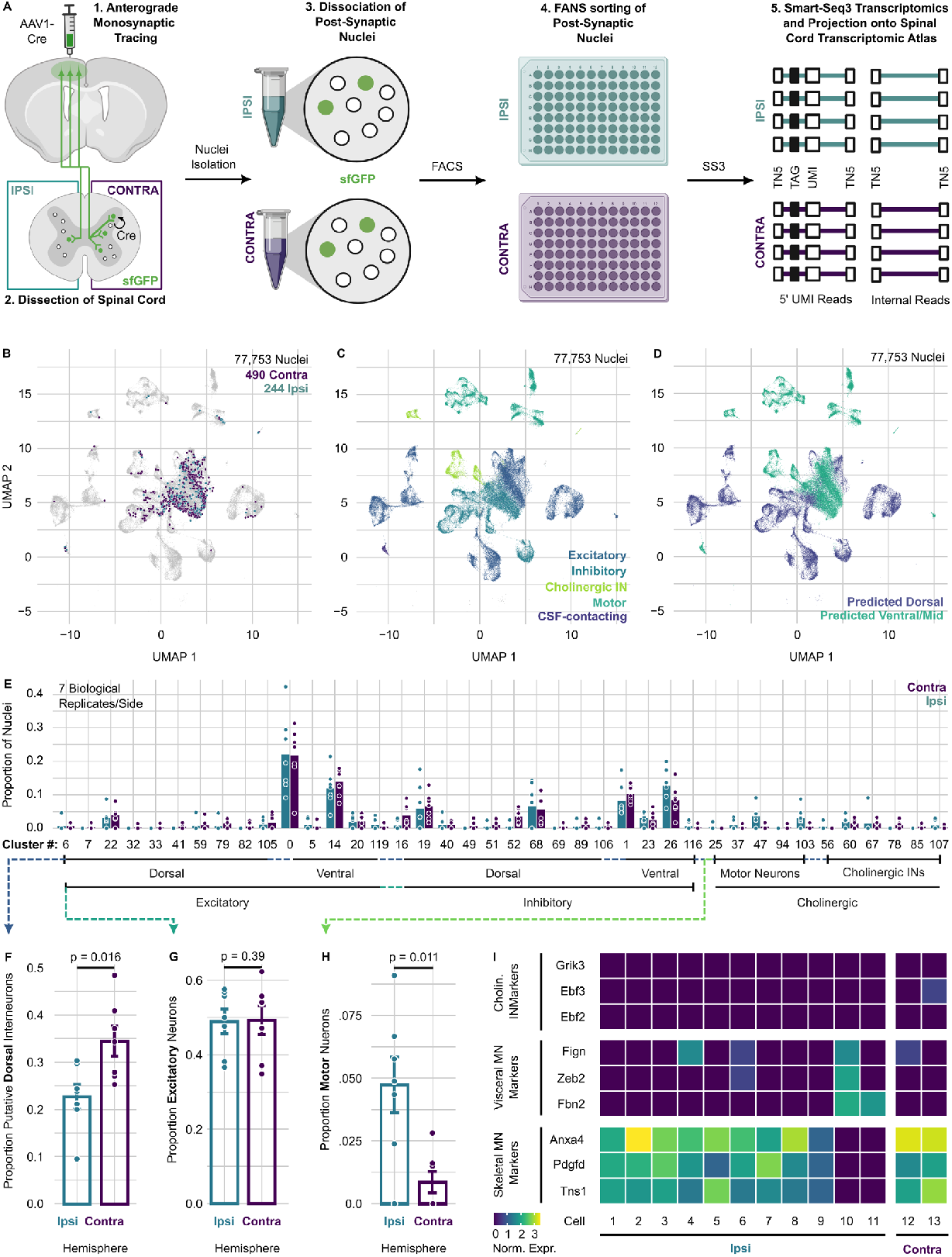
Comprehensive transcriptomic characterization of postsynaptic cells targeted by ipsilateral and contralateral CSN axons. **(A)** A schematic of the AAV1-Cre monosynaptic tracing experiment and sequencing of post-synaptic nuclei from both sides of the spinal cord (n = 7 pooled or unpooled biological replicates on both sides). **(B)** A projection of the sorted post-synaptic nuclei onto an integrated UMAP of 77,753 spinal cord neuronal nuclei. **(C)** A UMAP of coarse-grained clusters in the entire data set. **(D)** Inferred dorsal or middle-to-ventral-located clusters using the Russ *et al*. 2021 spinal cord atlas labels. **(E)** Bar plots of all neuronal clusters to which sorted post-synaptic nuclei are projected, expressed as a proportion of all sorted nuclei from either side and arranged by putative location and neurotransmitter type. **(F)** The fraction of inferred dorsal and middle-to-ventral localization among post-synaptic nuclei from either side of the spinal cord (p = 0.016, n = 7 biological replicates each, unpaired two-sided t-test). **(G)** The fraction of excitatory post-synaptic nuclei on each side of the spinal cord of each biological replicate (p = 0.39, n = 7 biological replicates each, unpaired two-sided t-test) **(H)**. The fraction of MN nuclei from either side of the spinal cord (p = 0.011, n = 7 each, unpaired two-sided t-test). **(I)** Among the 13 neurons that projected to MN clusters, a heat map showing the expression of markers from Alkaslasi *et al*. 2021 of skeletal MNs, visceral MNs, and cholinergic interneurons.

We note that this technique also labeled endothelial (5.8%) and oligodendrocyte (4.0%) nuclei, as evidenced by tagged nuclei in the spinal tissue periphery and dorsal funiculus as well as nuclei expressing their associated markers **(Fig S1E−G)**. The endothelial cell labeling could result from viral leakage into the cerebrospinal fluid, so *Pkd2l1-*expressing cerebrospinal fluid-contacting neurons were also excluded from downstream statistical analyses. Previous monosynaptic rabies tracing studies have documented viral transfer between oligodendrocyte precursor cells and neurons, which may explain the labeling of oligodendrocytes.^51^ After quality control, the resultant neuronal data set comprised 490 contralateral and 244 ipsilateral nuclei.

The sequenced post-synaptic nuclei provided information about neurons that received corticospinal input but did not provide information about the transcriptomic clusters that lack such input and were unrepresented in the data. We therefore generated an snRNA-Seq spinal cord data set comprising 73,120 sequenced nuclei to provide this transcriptomic context (to be described in detail elsewhere, Y.G, Z.Y., S.W.W., H.Z., and B.T., in preparation). We combined this data set with published spinal cord data sets containing 52,623 cells and nuclei and another containing 16,042 cholinergic-enriched nuclei **(Fig S2A−C)**.^52–59^ The snRNA-seq transcriptomes from our post-synaptically labeled nuclei were then projected onto this assembled atlas **(Fig 2B−E)**. One of the published analyses with which we merged our data annotated their clusters according to their position in the spinal cord **(Fig S2D)**.^53^ We used these annotations to label each cluster as likely located in the dorsal or likely mid-to-ventral laminae **(Fig S2E, Fig 2D)**. In line with our spatial analysis of post-synaptic neurons and axons **(Fig 1F−G, Fig 1J−K)**, there were fewer ipsilateral nuclei in clusters predicted to come from the dorsal horn relative to the contralateral nuclei (p = 0.016, n = 7, unpaired two-sided t-test) **(Fig 2F)**. There was no significant difference in glutamatergic and GABAergic/glycinergic nuclei proportions between the two sides (p = 0.39, n = 7, unpaired two-sided t-test) **(Fig 2G)**.

Motor neuron (MN) clusters were significantly over-represented in ipsilateral (4.9 ± 1.1%) compared to contralateral (0.9 ± 0.4%) post-synaptic nuclei, representing a 5.7-fold enrichment (p = 0.011, n = 7, unpaired two-sided t-test) **(Fig 2H)**. Examining markers^52^ of cholinergic interneurons (*Grik3, Ebf2, Ebf3*), visceral MNs (*Fign, Zeb2, Fbn2*), and skeletal MNs (*Anxa4, Pdgfd, Tns1*), most putative MN nuclei expressed skeletal MN markers **(Fig 2I)**. Greater connectivity to MNs is further evidence that ipsilateral CSN axons project to neuronal populations involved in motor control.

### The ipsilateral axons of the CST innervate spinal motor neurons

Many studies in rodents have failed to show direct synaptic connections between CSNs and MNs after activity-dependent pruning.^60–65^ However, sparse connections have been described, most recently in mice using monosynaptic rabies tracing from spinal MNs and direct visualization of CST synapses onto MNs.^25,64,66–69^ It is nonetheless essential to rule out potential avenues for non-specific labeling of MNs due to mistaken clustering,^70^ local synaptic leakage of the virus,^43^ or systemic leakage of AAV1-Cre to the neuromuscular junction.^71,72^

To do this, we labeled choline acetyltransferase (ChAT) in the spinal cords from Ai14 mice that received unilateral AAV1-Cre injections in the CFA. We enumerated tdT(+) ChAT(-) neurons throughout the cervical spinal cord and tdT(+) ChAT(+) neurons within MN pools on either side in cleared, whole cervical spinal cords **(Fig 3A−B, 3E−F)**.^73^

**Fig 3:**
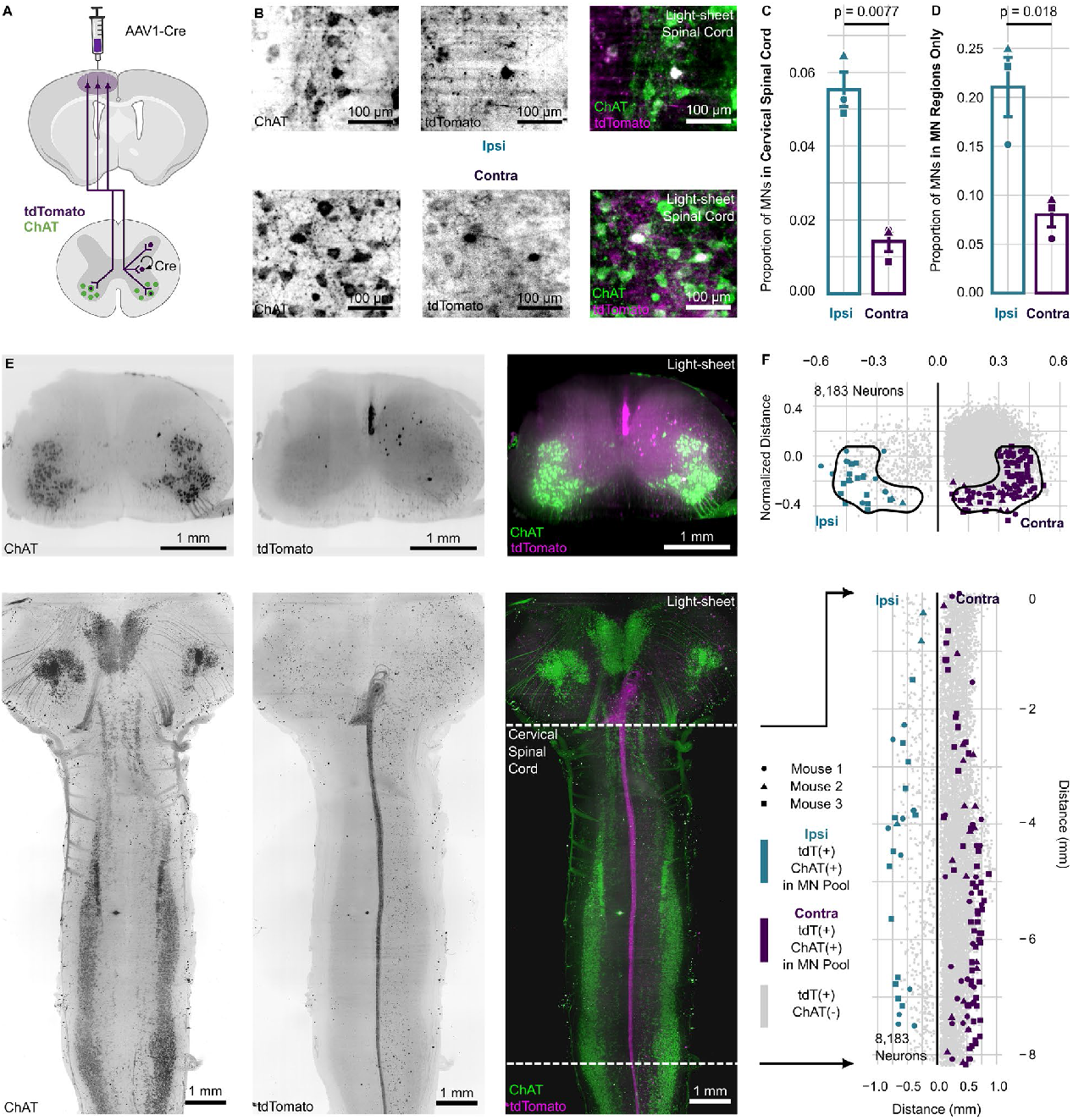
The ipsilateral axons of the CST innervate spinal motor neurons. **(A)** A schematic of the experiment (n = 3), in which the spinal cord tissue from the AAV1-Cre monosynaptic tracing in Fig 1 was stained for ChAT. **(B)** Images of antibody-stained MN pools and native post-synaptic tdT fluorescence, each a maximum-intensity projection of 100 μm. **(C)** A bar plot showing the share of ChAT(+) post-synaptic neurons as a proportion of tdT(+) post-synaptic neurons on their respective side, with the ipsilateral side in teal and the contralateral side in purple (p = 0.0077, n = 3, paired two-sided t-test). **(D)** A bar plot showing the same proportions as (C) but restricted only to the MN pool regions designated by the black outlines in (F) (p = 0.018, n = 3, paired two-sided t-test). **(E)** Images of the cleared spinal cords. On top are horizontal maximum-intensity z-projections 100 μm thick, and on the bottom are maximum-intensity z-projections from a coronal perspective. The analysis was conducted on regions inside the dotted lines. The brightness of representative images was adjusted in each channel so that tissue was visible via autofluorescence. **(F)** Dot plots of the locations of 8,183 post-synaptic neurons on a coronal plane. On the top is the same data on a horizontal plane in coordinates normalized to the width and height of the spinal cord grey matter, with MN pool regions used for (D) outlined in black. On the bottom is the same data plotted coronally. tdT(+) are plotted in gray, ipsilateral ChAT(+) tdT(+) in teal, and contralateral ChAT(+) tdT(+) in purple.

The number of ipsilateral and contralateral post-synaptic MNs as a fraction of total post-synaptic neurons on their respective side was similar to that in our transcriptomic data **(Fig 3C)**. Of contralateral tdT(+) neurons, 1.4 ± 0.3% were ChAT(+) and in MN pools, similar to 0.9 ± 0.4% in our transcriptomic data. The ipsilateral tdT(+) neurons were 5.5 ± 0.5% ChAT(+) and located in MN pools, in line with the 4.9 ± 1.1% in our transcriptomic data. This amounted to a 3.9-fold difference in ipsilateral/contralateral labeling preference (p = 0.0077, n = 3, paired two-sided t-test), comparable to the previously calculated 5.7-fold difference in our transcriptomic data **(Fig 2H)**. Replicating our findings *in situ* shows that our results were not due to mistaken cluster assignments.

Another alternate explanation for the increase in MN labeling on the ipsilateral side of the spinal cord is the local non-specific leakage of AAV1-Cre. If this were the case, we would expect equal proportions of MNs to be labeled in the immediate vicinity of MN pools because the rate of leakage ought to be the same on both sides within a local area of tissue. Even within a restricted regional analysis **(Fig 3D, F)**, there was still a 2.6-fold higher proportion of MNs tagged on the ipsilateral side (p = 0.018, n = 3, paired two-sided t-test). While some degree of non-specific labeling likely occurs, the proportionally larger labeling of MNs on the ipsilateral side cannot be explained by local non-specific leakage of AAV1-cre.

Finally, 79.8 ± 3.8% of all post-synaptic MNs were on the contralateral side (p = 0.016, n = 3, one sample, two-sided t-test, μ = 0.5). This indicates that our results were not due to systemic leakage of the virus to the neuromuscular junction, which would result in similar absolute numbers of tagged MNs on either side.

### IP-CSNs are primarily bilaterally-projecting neurons

We next asked if the ipsilaterally terminating CSN axons arose from neurons that only project ipsilaterally or whether they were collaterals of bilaterally projecting neurons. The commissural crossing observed in our BDA staining **(Fig 1I)** suggests bilateral projections, as do instances of individual bilateral axons found in primates.^74^ However, the pyramidal tract of the mole serves as a valuable counterpoint: its corticospinal axons descend ipsilaterally and decussate *en masse* to the contralateral side at the level of the spinal cord, suggesting that CSNs in mice could similarly decussate entirely to the ipsilateral side.^75^

To approach this question, we turned to MAPseq, a technique in which a Sindbis virus library encoding a high diversity of barcodes is injected into a population of neurons and used to reconstruct axonal projections. Infected neurons highly express and transport the barcode mRNA into their distal synapses at projection targets. Each neuron is uniquely labeled by random RNA barcodes, making individual barcodes correspond to a single neuron of origin. By sequencing the barcode mRNA from both injection and projection sites, it is possible to infer whether individual neurons project to multiple locations. To probe CSN projections, we injected the Sindbis virus library into the CFA, allowed it to express for 44 hours, and then dissected both sides of the cervical spinal cord and injection site for barcode sequencing **(Fig 4A, S3A)**. We also harvested and sequenced barcodes from the contralateral cortex as a positive control because many cortical neurons project to their opposite cortical hemisphere but not to the spinal cord. Olfactory bulbs served as negative controls since they receive no cortical input.^76^

**Fig 4:**
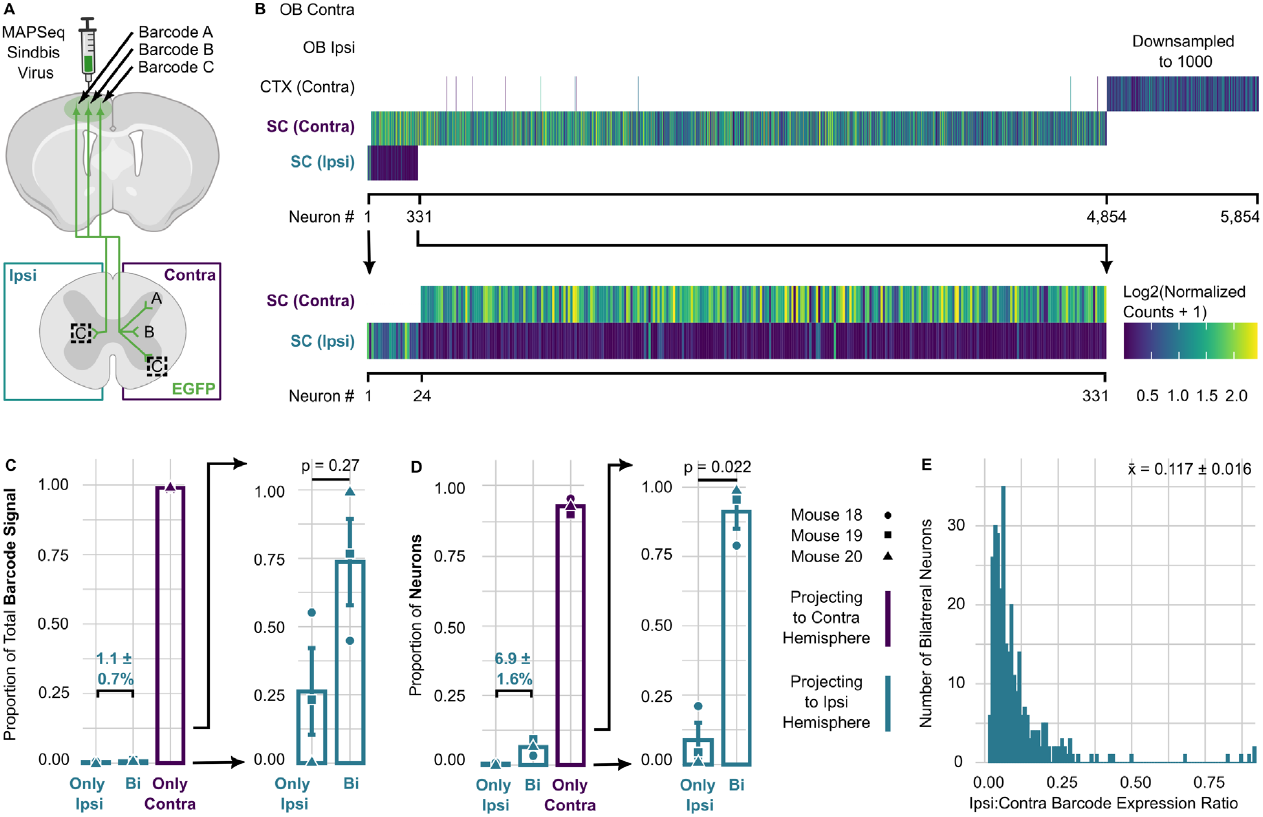
IP-CSNs are primarily bilaterally-projecting neurons. **(A)** A schematic of the MAPseq experiment (n = 3). **(B)** A heatmap displaying the 5,854 sequenced barcodes from the experiment, each representing an individual neuron. Each neuron is represented by a column of stripes, with the color corresponding to the log2-transformed normalized counts for that barcode. Each horizontal row represents a dissected region. The row representing the positive control commissural cortical projections is downsampled to 1000 neurons. Below, the same plot is displayed but magnified to the barcodes detected in the ipsilateral spinal cord. **(C)** Two bar plots show the distribution of the total normalized corticospinal barcode signal detected in the spinal cord. In the first chart, the share of barcode signal detected only on the ipsilateral side is in teal, and the share on the contralateral side in purple. The following bar plot examines only the proportions of the barcode signal detected on the ipsilateral side or both sides (p = 0.27, n = 3, one-sample, two-sided t-test, μ = 0.5). **(D)** Two bar plots show the distribution of neurons detected in the spinal cord. In the first chart, the share of neurons detected only on the ipsilateral side is in teal, and the share on the contralateral side is in purple. The following bar plot examines only the proportions of neurons detected on the ipsilateral side or both sides (p = 0.022, n = 3, one-sample two-sided t-test, μ = 0.5). **(E)** A histogram of the expression ratio for each unique barcode detected on both sides of the spinal cord (ipsilateral: contralateral) (p = 0.0017, n = 3, one-sample two-tailed t-test, μ = 0.5).

The ipsilateral side of the spinal cord contributed 1.1 ± 0.7% of total normalized CSN barcodes **(Fig 4B−C)**, similar to the 2.4 ± 0.9% of axons we tabulated with BDA tracing. After quality control, we detected 4,854 unique barcodes in the cervical spinal cord, which correspond to individual neurons. Of all unique barcodes in the cervical spinal cord, 6.9 ± 1.6% were present on the ipsilateral side **(Fig 4B, 4D)**, suggesting that approximately 6.9% of corticospinal neurons sent detectable axons to the ipsilateral spinal cord. Among these unique ipsilateral barcodes, 91.2 ± 6.2% (p = 0.022, n = 3, one-sample two-tailed t-test, μ = 0.5) were present on both sides of the spinal cord **(Fig 4B, 4D)**. This suggests that most IP-CSNs send axons to both sides of the spinal cord. For those barcodes detected on both sides, the average barcode expression on the ipsilateral side was 11.7 ± 1.6% of their corresponding contralateral expression (p = 0.0017, n = 3, one-sample two-tailed t-test, μ = 0.5) **(Fig 4E)**. This implies that ipsilaterally terminating axons are largely minor collaterals of bilaterally-projecting neurons that still predominantly terminate on the contralateral side. In contrast, the 8.8% ± 6.2% of ipsilateral barcodes found only on the ipsilateral side had relatively high barcode expression such that they made up 26.3 ± 15.8% of the total ipsilateral barcode reads (p = 0.27, n = 3, one-sample two-tailed t-test, μ = 0.5) **(Fig 4C)**.

Taken together, these data are consistent with ipsilaterally terminating corticospinal axons comprising a mixture of two groups of neurons: a minor population that projects completely or near-completely ipsilaterally and a much larger population that is bifurcated and projects collaterals to the ipsilateral side while mostly terminating contralaterally. These data also reveal that while the ipsilaterally terminating axons are sparse, they represent a population of neurons substantially more numerous than their small number of axons suggests.

### The cell bodies of IP-CSNs have a specific cortical distribution

Analyzing the distribution of cortical neurons across their laminar surface has proven to be a valuable approach to understanding their organization and function.^77–80^ If the sparse ipsilaterally projecting axons constitute a substantial fraction of CSNs, we wondered where they were distributed across the motor cortex. For this reason, we next mapped the cortical topography of IP-CSNs using retrograde tracing.

We injected two retrograde AAVs^81^ expressing either mEmerald or mCherry-tagged TOMM20, which label cell bodies, unilaterally into C5−C6 of opposite sides of the spinal cord, placing injections such that they did not cross the spinal cord midline **(Fig 5A−C)**. After 21-24 days of expression, we harvested the cortices and spinal cords of injected mice and imaged them with CUBIC tissue clearing on a lightsheet microscope. Quantification of 6,834 mEmerald(+) and mCherry(+) neurons in the cortex showed 4.8 ± 0.3% were located ipsilateral to their injection site **(Fig 5E)**, similar to the 6.9 ± 1.6% figure found using MAPseq. Only 36.6 ± 8.6% of IP-CSNs were labeled with both fluorophores (p = 0.26, n = 3, one-sample two-tailed t-test, μ = 0.5), compared with the 91.2 ± 6.2% found with MAPseq **(Fig 5E−F)**. However, this number is difficult to compare to our MAPseq data because double-labeling percentages are the coincidence of two separate virus injections and their incomplete labeling efficiencies.^82^ For this reason, the MAPseq data are likely the more accurate measure of the proportion of bilateral neurons. Contralateral neurons were distributed among the Rostral Forelimb Area (RFA), the CFA, and a caudo-lateral cluster, which has been referred to as S2 **(Fig 5D)**.^46,83^ The IP-CSNs were nearly absent from S2 (p = 0.0056, n = 3, Bonferroni-corrected one-sample two-tailed t-test, μ = 0.5) **(Fig 5D)**, with double-labeled IP-CSNs being entirely absent (p = 0.030, n = 3, Bonferroni-corrected one-sample two-tailed t-test, μ = 0.5) **(Fig 5G−H)**.

**Fig 5:**
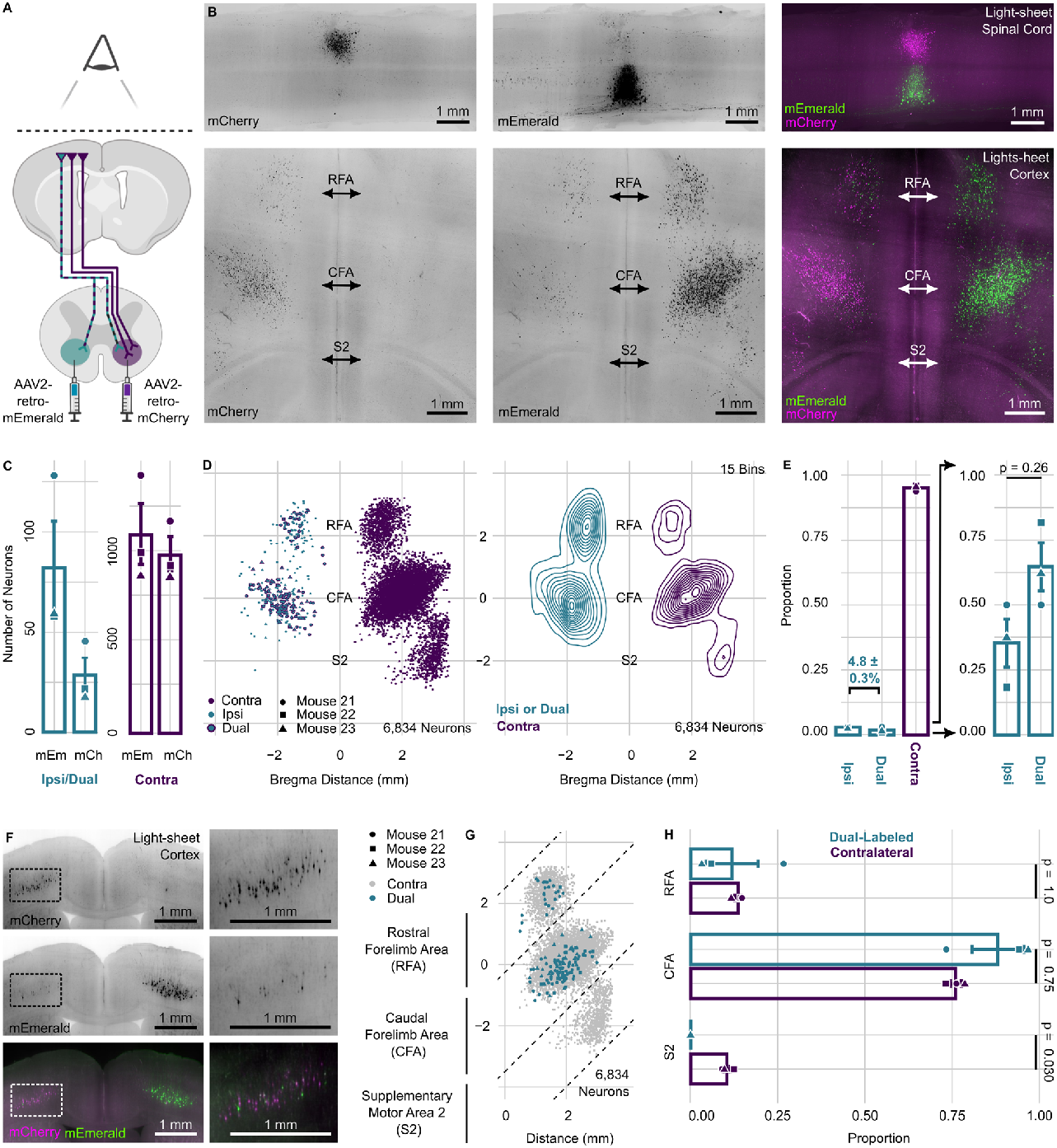
The cell bodies of IP-CSNs have a specific cortical distribution. **(A)** A schematic of the experiment (n = 3). **(B)** Representative maximum intensity z-projections of native mCherry and mEmerald fluorescence from cleared injection site tissue and motor cortices on the coronal and horizontal plane, respectively. The brightness of representative images was adjusted in each channel so that tissue was visible via autofluorescence. **(C)** Bar plots representing the number of annotated neurons by fluorophore and laterality plotted on a horizontal plane, with ipsilateral in teal and contralateral in purple. **(D)** First, a dot plot of 6,834 annotated neurons on a horizontal plane with ipsilateral neurons in teal, contralateral in purple, and double-labeled neurons in teal outlined with purple. Second, a contour plot of the same data (15 bins). **(E)** First, bar plots indicate the ratio of contralateral neurons in purple and ipsilateral and dual-labeled ipsilateral neurons in teal. The second bar plot examines only ipsilateral neurons labeled with a single fluorophore and those labeled with both (p = 0.26, n = 3, one-sample two-sided t-test, μ = 0.5). **(F)** Representative images of native mCherry and mEmeraled-labeled CSNs in the cortex on a coronal plane. **(G)** A dot plot of labeled neurons on a horizontal plane, with ipsilateral neurons in teal and contralateral neurons in gray. The dotted lines indicate segmentation of the three motor cortical regions. **(H)** Bar plots compare the ratio of double-labeled and contralateral neurons in each cortical region (S2: p = 0.030, n = 3, Bonferroni-corrected one-sample two-sided t-test, μ = 0.5).

IP-CSNs are present in the two motor-associated regions^9,84,85^ but absent from S2, a region only associated with sensory roles.^46,86^ These data show that ipsilateral corticospinal axons are not just a feature of the CST, but originate from separate neuronal populations with specific cortical distributions.

### IP-CSNs bear an embryonic-like molecular signature related to neural regeneration

Since IP-CSNs have distinct projection targets, a distinct relationship with the midline boundary, and exist in distinct cortical regions, we tested if this host of differences might manifest in distinct gene expression. To do so, we used the viral implementation of Translating Ribosome Affinity Purification (vTRAP) to examine the molecular properties of these cells.^87^

We injected spinal cord segments C5−C6 unilaterally with a retrograde AAV expressing the EGFP-L10a vTRAP transgene and allowed the virus to express for 25–27 days **(Fig 6A, Fig S4A−D)**. EGFP-tagged polysomes were then immunoprecipitated (IP) from homogenates of each cortical hemisphere using anti-EGFP antibodies. Polysome-bound mRNA was then isolated and analyzed by RNA-seq. PCA analysis showed TRAP IP samples clustered similarly compared to their whole tissue input samples **(Fig S4E)**. Gene sets of both canonical markers of Layer 5b pyramidal neurons (p = 0.31, n = 5, CAMERA gene set test, seven genes) and markers derived from a bacTRAP line labeling Layer 5b neurons^88^ (p = 0.31, n = 5, CAMERA gene set test, 17 genes) were similarly expressed in both populations **(Fig S4F)**. This is consistent with equivalently efficient pulldowns.

**Figure 6:**
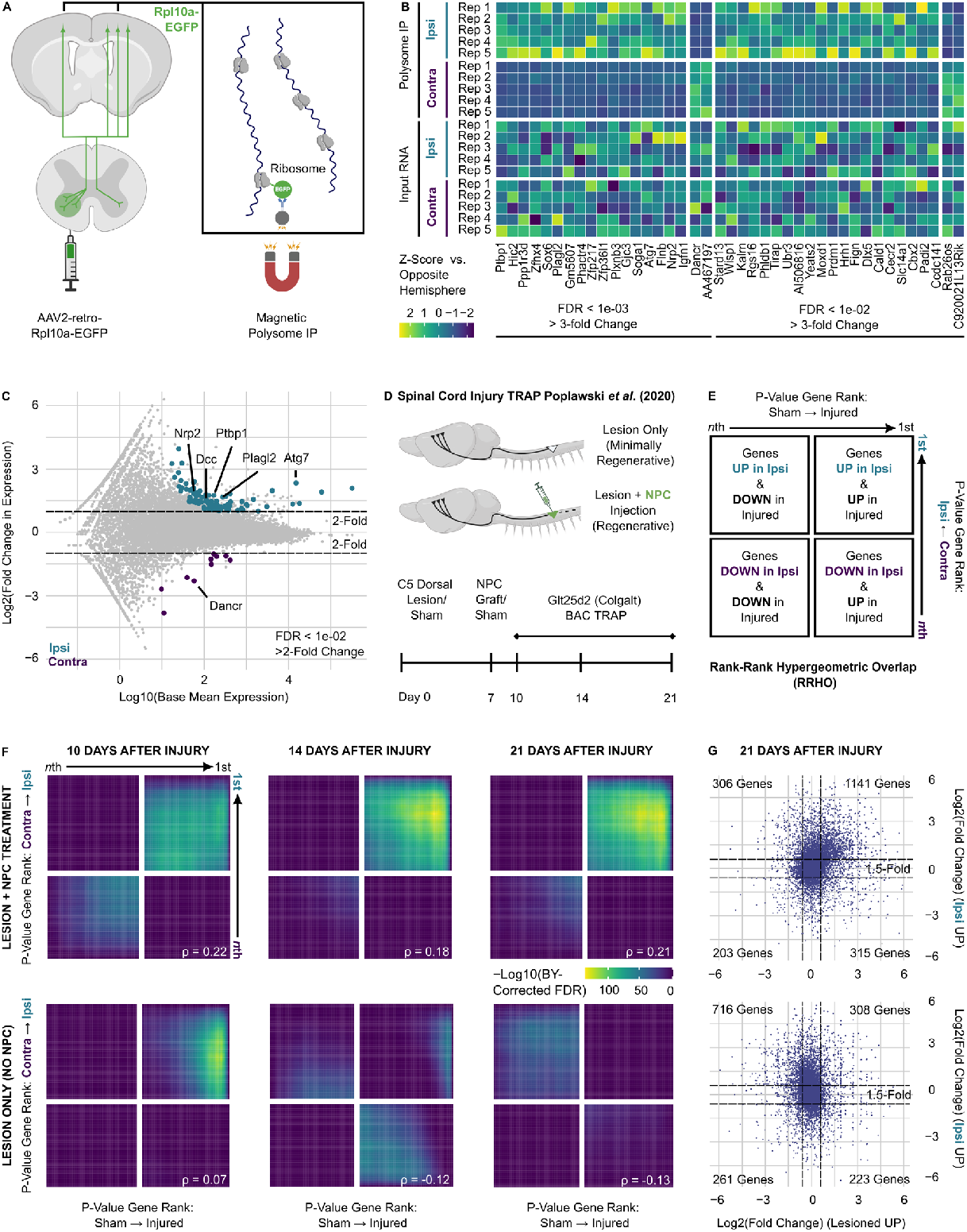
IP-CSNs bear an embryonic-like molecular signature related to neural regeneration. **(A)** A schematic of the viral TRAP experiment (n = 5). **(B)** A heatmap of z-scores for significantly enriched genes (DESeq2 FDR < 0.01 and >3-fold Change) from ipsilateral neurons in teal and contralateral neurons in purple. Z-scores were calculated separately between all precipitated polysome samples and all unprecipitated input RNA samples. **(C)** An MA plot of Log2(fold-change) differences in expression between ipsilateral and contralateral populations. Genes with an FDR < 0.01 and >2-Fold Change are colored teal if upregulated in ipsilateral neurons and purple if in contralateral. Genes discussed in the main text are indicated. **(D)** A schematic of Poplawski *et al*. 2020’s bacTRAP experiments (n = 2–3) in injured CSNs. **(E)** A schematic of Rank-Rank Hypergeometric Overlap (RRHO) analysis, as performed in subsequent panels. **(F)** RRHO analysis comparing the differential expression of ipsilateral vs. contralateral CSN populations with those of Poplawski *et al*. (2020). The top row of plots compares our vTRAP differential expression data with that of NPC-treated, regenerating corticospinal neurons throughout their recovery and the bottom to the untreated, minimally regenerating corticospinal neurons. Positive Spearman’s ρ values indicate differential expression concordant with that of IP-CSNs. FDR values are expressed using the Benjamini-Yekutieli method for multiple comparison correction. **(G)** Scatter plots of Log(2-fold) changes compare the vTRAP differential expression data of our own ipsilateral vs. contralateral CSNs to the 21-day post-injury time point samples of Poplawski *et al*. (2020). The top panel compares the present neurons of the study to the NPC-treated, regenerating neurons, and the bottom panel compares the untreated neurons.

Using the DESeq2 pipeline (FDR < 0.05), we found 585 genes were differentially expressed between the two populations **(Fig 6B−C)**. Notably, *Dcc*, a gene whose knockdown globally disrupts midline crossing,^28,30^ was expressed 2.8-fold higher in IP-CSNs (FDR = 0.0047). We performed gene set enrichment analysis (GSEA) with Gene Ontology (GO) terms for biological processes on genes enriched in IP-CSNs **(Fig S5A)**. Developmental processes (e.g., forebrain development, telencephalon development, CNS neuron development) dominated the top terms in this analysis.

We noticed that among the 19 most significantly differentially expressed genes (FDR < 0.001, >3-fold change), several have been independently linked to neuronal degeneration and regeneration. *Ptbp1* (3.4-fold higher in ipsilateral, FDR = 6.7e-14) knockdown impairs neural regeneration in peripheral sensory neurons,^89^ ribosome-associated lncRNA *Dancr*^90^ (4.9-fold higher in contralateral, FDR = 3.6e-10) increases murine spinal cord neuron apoptosis and spinal cord inflammation after injury,^91^ Nrp2 (3.9-fold higher in ipsilateral, FDR = 7.1e-4) antibody blockade reduces murine corneal nerve regeneration,^92^ *Plagl2* (3.2-fold higher in ipsilateral, FDR = 5.9e-6) induction increased the regenerative potential of aged murine hippocampal neural stem cells,^93^ and *Atg7* (5.0-fold higher in ipsilateral, FDR = 4.3e-4) conditional knockdown in murine Purkinje neurons causes axonal degeneration.^94^

A study using TRAP found that following CST lesions, adult CSNs exhibited molecular similarities to embryonic CSNs, which persisted long-term when neural precursor cells (NPCs) were injected into the injury site. The authors reasoned that the persistence of this embryonic-like molecular signature in the NPC-treated mice allowed the CSNs to regenerate through and beyond the lesion **(Fig 6D)**.^95,96^ Several of the genes differentially expressed in the ipsilateral neurons corresponded to those found in the regenerative signature. To examine if these single-gene concordances were part of broader similarities between the data sets, we adopted the threshold-free transcriptome-wide analysis method used by Poplawski *et al*. (2020).^95^ Rank-rank hypergeometric overlap (RRHO) is a non-parametric and threshold-free method of comparing differential expression experiments.^97,98^ Ranked p-values are plotted on the x- and y-axis, and the resultant scatter plot is displayed as a heatmap that can be read to determine the strength and pattern of correlation across the entirety of differentially expressed genes between two experiments **(Fig 6E)**. When we compared the differential expression between ipsilateral and contralateral CSNs to each of the data sets from Poplawski *et al*. (2020), IP-CSN enriched genes showed a strong correlation with the regenerative molecular signatures (ρ = 0.07−0.22, maximum Benjamini-Yekutieli-corrected FDR = 1e-136.3–1e-88.0) and an anticorrelation with the non-regenerative signatures (ρ = -0.13– -0.12, maximum Benjamini-Yekutieli-corrected FDR = 1e-57.7–1e-48.2) **(Fig 6F)**. For example, 1,141 genes were commonly upregulated (>1.5 fold) in both IP-CSNs and the NPC-treated regenerative mice at 21 days, in contrast to only 308 genes co-upregulated in the untreated non-regenerative mice at the same time point **(Fig 6G)**. The RRHO plots indicated a set of genes concordantly upregulated but few concordantly downregulated. Repeating GSEA analysis on these concordantly upregulated genes across the regeneration-associated time points similarly yielded terms associated with developmental processes **(Fig S5B-F)**.

Finally, employing the same gene expression data set^99^ of developing CSNs used by Poplawski *et al*. (2020)^95^ **(Fig S6A−B)**, we likewise found an anticorrelation between genes differentially expressed in IP-CSNs and the molecular changes that occur from their embryonic to post-natal transcriptomes (ρ = -0.17– -0.12, maximum Benjamini-Yekutieli-corrected FDR = 1e-28.6–1e-15.4) **(Fig S6C)**.

Taken with the enrichment of development-associated GO terms, this shows that the molecular characteristics of IP-CSNs also resemble those of their embryonic form.

### IP-CSNs are spontaneously regenerative

In addition to IP-CSNs exhibiting a regenerative-like molecular signature, several of their anatomical properties are also confluent with those of regenerating^100^ corticospinal neurons. These include selective innervation of motor-associated spinal populations in NPC-treated animals,101 bilateral extension of regenerating axons into the spinal gray matter in pro-regenerative cortical *Pten* knockdowns,102 and the observation that ipsilaterally-terminating axons of the ventral CST are more regenerative than axons from the contralateral dorsal funiculus in a model of cervical re-innervation following thoracic hemisection.103 We therefore asked if IP-CSNs might represent an inherently more regenerative subpopulation.

We elected to use a model in which lumbar projecting CSNs are severed via a T8 dorsal hemisection, resulting in increased innervation of the cervical spinal cord and concomitant functional recovery. This paradigm has been extensively replicated and models naturalistic recovery from injury rather than aiming to enhance regeneration artificially.^103–109^

The experimental design was based on published reports examining this phenomenon in rats **(Fig 7A)**.^106,108^ The ipsilateral and contralateral CSNs were labeled by unilaterally injecting retrograde AAVs expressing nuclear-localized mGreenLantern (mGL) on one side of the spinal cord at segments L2–L4 and mScarlet (mSc) on the opposite side **(Fig S7A−B)**.^82^ Ten days after AAV injections, we performed a dorsal hemisection (DHX) or sham surgery at segment T8 **(Fig S7C)**. After an additional 21 days of recovery, we injected Fluoro-Gold (FG) into the cervical spinal cord at segments C3–C4 on both sides **(Fig S7D)**. We then harvested tissue and imaged every third serial section using an epifluorescence microscope. Consistent with prior studies,^103–105^ we normalized the proportions of FG(+) lumbar-projecting CSNs to the total number of FG(+) neurons in the hindlimb field of the motor cortex to compensate for the varying efficiencies of FG injection. If IP-CSNs are more regenerative, a greater proportion of these would contribute to the increased innervation of the cervical level and thus be co-labeled with FG.

**Fig 7:**
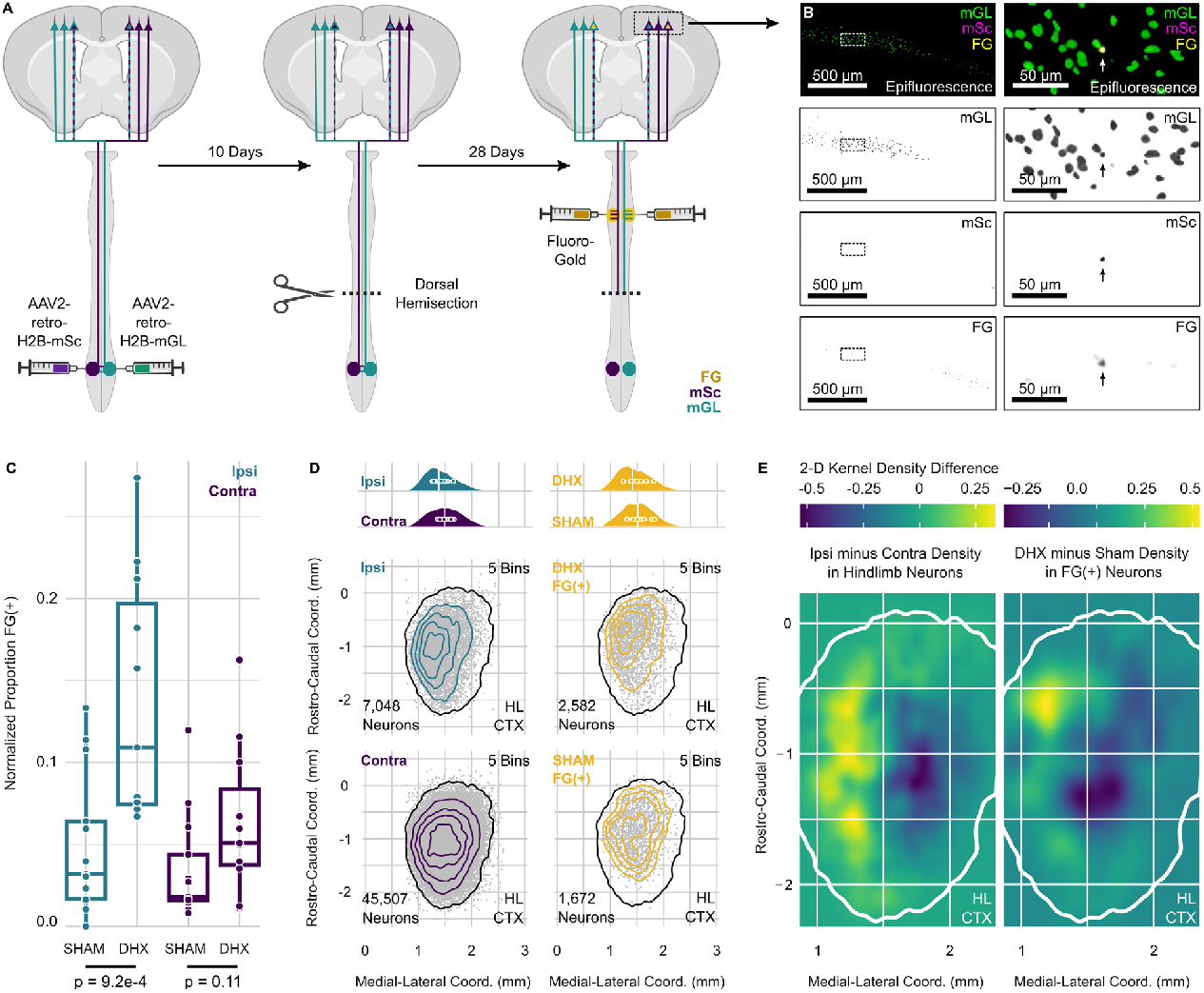
IP-CSNs are spontaneously regenerative. **(A)** A schematic of the experiment (n = 11 DHX, 13 sham). **(B)** Fluorescent images of a 50 μm coronal slice of hindlimb field, with lumbar-projecting neurons with native mGreenLantern expression in green, lumbar-projecting neurons with native mScarlet expression in purple, and cervical projecting neurons with native FG fluorescence in yellow. Representative images were thresholded in each channel to match the values used for analysis. **(C)** Bar plots showing the proportions of IP-CSNs (p = 9.2e-4, n = 11 DHX, 13 sham, two-sided unpaired Wilcoxon signed-rank test) and non-IP-CSNs (p = 0.11, n = 11 DHX, 13 sham, two-sided unpaired Wilcoxon signed-rank test) double-labeled with Fluoro-Gold in both the sham and dorsal hemisection (DHX) groups. Proportions were normalized to the total number of FG(+) neurons in each hemisphere to account for injection variability. **(D)** Dot plots of the locations of 52,555 neurons on a horizontal plane. The black outline designates the hindlimb cortex (99% of lumbar-projecting neurons). Density plots (5 bins) of their distribution are overlaid, with contralateral neurons in purple, ipsilateral in teal, and FG(+) in gold. Above are ridgeline plots showing the medial-lateral distribution of the neurons, comparing ipsilateral to contralateral and FG(+) sham to FG(+) DHX neurons. **(E)** On the left is a heat map showing differences in 2-dimensional kernel density estimation between neurons with ipsilateral projections and those without. Greater values indicate a greater relative density of neurons with ipsilateral projections in that area. On the right is a heatmap showing the same differences between FG(+) neurons in hemisected and sham mice. Here, greater values indicate greater relative density in FG(+) neurons of the hemisected mice relative to the sham. In both, the same hindlimb cortex contour from (D) is plotted in white.

We analyzed 52,555 CSNs expressing the retrograde AAVs, with similar numbers tagged in the sham and DHX groups (p = 0.89 mGL and 0.43 mSc, n = 11 DHX, 13 sham, two-sided unpaired t-test) **(Fig 7B, Fig S7E)**. The total number of FG(+) neurons in the hindlimb field was also similar between groups (p = 0.33, n = 11 DHX, 13 sham, two-sided unpaired t-test) **(Fig S7F−G)**. Manual counting of labeled nuclei found 13.5 ± 0.8% of neurons in the ipsilateral cortical hemisphere. Normalized to FG injection efficiency, 3.5 ± 0.9% of contralateral CSNs were FG(+) in the sham group and 6.4 ± 0.7% in the DHX group (p = 0.11, n = 11 DHX, 13 sham, unpaired two-sided Wilcoxon signed-rank test) **(Fig 7C)**. Among IP-CSNs, 5.0 ± 1.2% were FG(+) in the sham group, compared to 14.1 ± 1.1% in the hemisected mice (p= 9.2e-4, n = 11 DHX, 13 sham, unpaired two-sided Wilcoxon signed-rank test) **(Fig 7C)**. These conclusions do not depend on the stringency of thresholds used to annotate FG(+) neurons or IP-CSNs **(Fig S7H)**.

The increase in co-labeling from ipsilaterally-projecting CSNs was also reflected in a shift in the distribution of FG(+) neurons toward the regions of the cortex occupied by IP-CSNs **(Fig 7D−E)**. This rostro-lateral distribution of double-labeled neurons upon hemisection toward the region in which we observe ipsilaterally-projecting CSNs was also independently observed in prior studies of compensatory cervical innervation.^106^ Taken together, our data indicate that increased innervation by lumbar-projecting axons at the cervical level is disproportionately, if not primarily, due to the sprouting of IP-CSNs.

## Discussion

The involute heterogeneity of nervous systems complicates our understanding of biological processes in the brain.^110,111^ Supposing disease-relevant phenomena are isolated to one class of cells, experiments searching for them in a heterogeneous population will diminish signal and risk missing patterns essential for discovery. To this end, the discovery of IP-CSNs as a distinct subpopulation of cortical afferents to the spinal cord stands to enhance the resolution of experiments across a wide range of investigations and may allow discoveries that would otherwise be mired in biological noise. Because traction into this class is only a matter of looking to the other side of the brain, the exquisite toolbox of mouse molecular neuroscience is available to those who wish to study them with a minimal technical barrier to entry.

Our findings suggest that focusing on IP-CSNs will be especially important for studies of spinal cord injury and cortical trauma that result in devastating paralysis.^112^ CSNs are among the least regenerative cell types in the nervous system,^96,113,114^ making detailed examination of the molecular mechanisms of injury response and their activity during recovery challenging to parse from experimental background. Anderson *et al*. (2022)^115^ write of the need to identify “recovery-organizing” populations of neurons to guide future investigations of recovery from spinal cord injury. We show that IP-CSNs share several features with regenerating CSNs, including bilaterality, projection to ventral (motor) spinal cord regions, direct connectivity to motor neurons, and a specific embryonic-like molecular signature. In addition, our results in a thoracic hemisection model provide direct evidence that these neurons are also spontaneously regenerative. The present results enable the specific examination of a subpopulation of spontaneously regenerative CSNs in relation to similar but minimally regenerative contralateral CSNs. Our results are paralleled by similar findings using transcriptomic data as a guide to discern and prioritize spinal cord neurons that are particularly responsive to injury.^116–118^

Much work in the fields of spinal cord injury and stroke already examines this population of neurons, if unintentionally.^25,119,120^ These experiments selectively damage one hemisphere of the cortex or pyramidal tract and study compensatory innervation (“sprouting”) from ipsilateral axons, reasoning that these axons represent an alternate circuit for descending information. In one sense, the well-established plasticity of ipsilateral corticospinal axons after injury supports our findings. However, this work was performed without awareness that such experiments might examine a specific class of neurons innately primed to compensate for injury. For example, Phase II clinical trials of NOGO-inhibiting antibodies are based in part on evidence from this experimental paradigm.^35,120^ Our results re-contextualize this large body of work and will be a critical experimental consideration for the field moving forward.

In a broader sense, in pursuing axons that depart from the prevailing pattern of corticospinal laterality, we have been led to a class of CSNs that are exceptional in many other respects. One coherent framework for organizing these observations is development.^3^ Before activity-dependent pruning, murine CSNs are more bilateral,^121^ project directly to motor neurons,^61,64^ and have greater regenerative capacity.^114,122,123^ IP-CSNs retain these juvenile characteristics into adulthood, including a molecular similarity to their juvenile form. This raises the possibility that other neuronal cell types might harbor subpopulations that retain juvenile transcriptional programs and anatomical characteristics into adulthood. Recent comprehensive single-nucleus sequencing projects have provided an invaluable portrait of the molecular landscape of mammalian nervous systems, but determining how to organize this complex and continuous heterogeneity in a biologically informed manner is an ongoing challenge.^124–128^ The retention of juvenile transcriptional programs and anatomical characteristics may be a broadly applicable principle for organizing the molecular and anatomical heterogeneity of adult nervous systems.

### Limitations

Ipsilateral corticospinal axons have been described in primates, with substantial termination in Lamina VIII.^20,21,74,129,130^ Comparing these data directly to our own is challenging because primate CSTs are more ventrally oriented at baseline.^1,5,131^ Our final experiment examines lumbar projecting corticospinal neurons, a distinct population of CSNs.^132^ However, there are similar numbers of ipsilateral CSN axons in the lumbar cord with similar ventral projection patterns.^46^ Given the lack of established experimental paradigms examining CSN regeneration in the cervical spinal cord, it was infeasible to approach the question without making this trade-off.

## Materials and Methods

Preprint formatting based on a publicly available template (https://github.com/chrelli/bioRxiv-word-template).

### I: Resource Availability

#### Lead Contact

Further information and requests for resources and reagents should be directed to and will be fulfilled by the lead contact, Eric Schmidt.

#### Materials Availability

- Plasmids generated in this study will be deposited to Addgene.

#### Data and Code Availability

- RNA-Seq data sets generated in this publication will be deposited at GEO. Accession numbers will be listed in the key resources table. Microscopy data associated with this publication is available from the lead contact upon request.
- Analysis code sufficient to reproduce the figures will be deposited on Zenodo.
- Any additional information required to reanalyze the data reported in this paper is available from the lead contact upon request.

### II: Experimental Models and Subject Details

All procedures were done in accordance with guidelines approved by the Institutional Animal Care and Use Committees at The Rockefeller University and the Institutional Animal Care and Use Committee of the Allen Institute for Brain Science, both in accordance with NIH guidelines.

#### Mice

All mice were experimentally naïve and purchased or were F1 progeny of mice purchased from The Jackson Laboratory. Mice were group housed in a 12hr/12hr light-dark cycle and provided *ad libitum* access to food and water. If the mice received spinal cord surgery, they were single-housed, as were any mice in cages with bite wounds or barbering from cage mates after surgery. Assignments of experimental groups were randomized among cages and sex if more than one cage of mice was used.

At euthanasia, mice were aged 15–16 weeks (2 M, 1 F) (anterograde tracing in Ai14 mice), 14–19 weeks (6 M, 5 F) (anterograde tracing in IN-TACT mice), 9–10 weeks (3 M, 2 F) (spinal cord snRNA-seq atlas), 41–45 weeks (1 M, 2 F) (MAPseq), 17–20 weeks (2 M, 1 F) (retrograde tracing of cortical distribution), 27–29 weeks (6 M, 9 F) (vTRAP), and 11–14 weeks (16 M, 7 F) (regeneration). Mice used for BDA tracing were >90 days old at the time of injection.^46^ Both male and female mice were used, and no significant sex differences were detected in the analysis; however, experiments were not powered to examine sex differences to keep these studies affordable. In some instances, female mice were under-represented because of exclusion criteria. Any mice <85% of starting weight were excluded from the study.

All mice were heterozygous Ai14 mice (The Jackson Laboratory, #007914; RRID:IMSR_JAX:007914) with a C57/B6 background and cross (The Jackson Laboratory, #000664; RRID:IMSR_JAX:000664), except for mice used for triple retrograde tracing after hemisection, which were wild-type C57/B6 mice, those used for Smart-seq3 sequencing, which were IN-TACT mice^48^ (The Jackson Laboratory, #021039; RRID:IMSR_JAX:021039), and the mice used for BDA tracing, which were wild-type Balb/C mice (The Jackson Laboratory, #000651; RRID:IMSR_JAX:000651).

## III: Method Details

### General Data Analysis and Visualization

Unless otherwise noted, data were analyzed with custom R^133^ (version 4.3.2, RRID:SCR_001905) script in RStudio^134^ (Build 494; RRID:SCR_000432), and figures were produced with the ggplot2 package (version 3.4.4; RRID:SCR_019186). Bar plots and dot plots were generated using the geom_bar, geom_errorbar, and geom_point functions; density plots were generated with the stat_density_2d package, box and whisker plots with the geom_boxplot function; histograms with the geom_histogram function, heat maps with the geom_tile or geom_raster function, and contour plots with the stat_density_2d function. Color schemes were generated with the Viridis package (version 0.6.4; RRID:SCR_016696). Figures were created using Inkscape (version 1.3, RRID:SCR_014479). Schematics were created using Biorender (RRID:SCR_018361).

### Representative Images

In most instances, the brightness of representative images was adjusted in each channel so that tissue was visible via autofluorescence. This is approximately the brightness and contrast used to annotate the images for pixel classification because signal/background discrimination was our primary concern. Figure 7 displays images with the exact signal thresholds employed in the analysis.

For the spinal cord, there is occasionally confusion as to the appropriate terminology for anatomical planes because the conventional mouse cerebral anatomical planes are technically incorrect with respect to their body orientation. This is to match the planes of mouse brains to those of humans, whose posture is upright and whose brains are rotated with respect to the body. We followed the convention used by the GENSAT project^110^ in which spinal cord planes are kept consistent with body plane orientation, i.e., “horizontal” slices proceed rostro-caudally and “coronal” planes proceed dorso-ventrally.

### General Surgical Procedures

Surgical procedures and viral injections were carried out under protocols approved by Rockefeller University IACUC. Mice were given DietGel® Recovery hydration gel cups (ClearH20) for one week before surgery and then acclimated for 24 hours in the surgery room. All injectables were heated to the mouse’s body temperature. The day before the procedure, they were given 1 mg/kg dexamethasone intraperitoneally (IP). Before the surgery, the mice were deeply anesthetized with 100 mg/kg ketamine and 10 mg/kg xylazine in normal saline injected IP, followed by 1 mL of normal saline for fluid support administered subcutaneously (SC), a second 1 mg/kg dose of dexamethasone in normal saline administered SC, and 6 mg/kg extended-release meloxicam administered SC. In the case of laminectomy, 3.25 mg/kg extended-release buprenorphine was additionally injected SC.

When the mice passed the toe pinch test, their eyes were covered with Paralube ophthalmic lubricant (Dechra), and hair around the area of the incision was shaved and removed with chemical hair removal (Nair Sensitive Formula). They were placed on oxygen (0.5 L/min) and heat (∼37° C) support for the remainder of the procedure. The skin was sterilized three times with ethanol washes. At most, 5 mg/kg bupivacaine was injected subcutaneously at the incision site for local anesthesia.

For intraspinal injections into the ventral horn of the cervical spinal cord, an incision was made from the base of the skull to the shoulder blades of the mouse, and the two layers of muscle above the spinal cord were opened and held apart with tissue separators. The muscle surrounding the spinal cord was carefully teased away with a dental scaler to reveal spinal vertebrae. Using the T2 vertebral spine as a landmark,^135^ vertebrae of interest were removed with blunt forceps (Fine Science Tools, 11223-20). The dura, but not the arachnoid and pia mater, was removed above the injection site with small scissors (Fine Science Tools, 15000-03) and forceps. After injections, three drops of 1.25 mg/mL bupivacaine were applied directly to the exposed spinal cord and surrounding tissue. Then, both layers of muscle were re-joined with three resorbable sutures (Vilet, V317), and two drops of bupivacaine were applied locally for pain relief. The wound was then sealed with tissue adhesive and secured with two staples.

For cranial injections, 0.1 mL of 1.25 mg/mL bupivacaine diluted in normal saline was injected along the scalpel incision site. Holes were drilled through the skull to the dura mater with a rotary tool (Dremel) and a 0.6 mm rounded carbide bur (Meisinger, HM1-006-RA), occasionally wet with sterile normal saline to prevent overheating. After injections, the wound was sealed with tissue adhesive.

Viruses were injected with an oil hydraulic micromanipulator (Narishige, MO-10) attached to a pulled glass PCR micropipette (Drummund, 5-000-1001-X) filled with mineral oil. The virus was injected over one minute on either side of the spinal cord, and the needle was allowed to rest for 5 minutes before being carefully withdrawn over 30 seconds.

After surgery, mice continued to receive oxygen support and heat support under one side of their cage until ambulatory, as well as recovery gels for the following two weeks or until euthanasia. On the first and second days after surgery, the mice received 0.5 mg/kg dexamethasone administered SC, and on the third day, they received 0.25 mg/kg before discontinuation. After three weeks, any remaining staples were removed under light isoflurane anesthesia. Mice were observed for locomotion irregularities, weight loss, grimacing, nesting behavior, and food and water consumption. Mice demonstrating visible deficits in limb control, decreases in weight, lack of nesting, or squinting/grimacing were excluded from the study and euthanized.

### General Light-sheet Procedures

#### CUBIC Tissue Clearing

Mice were euthanized with a ketamine and xylazine mixture in normal saline (200 mg/kg and 20 mg/kg, respectively). Immediately, the mice were perfused with 30 mL ice-cold PBS, followed by 30 mL ice-cold PBS with 4% paraformaldehyde. The brain and spinal cord were removed and placed overnight at 4° C in 4% paraformaldehyde before being washed three times for two hours in room-temperature PBS. Tissue was stored for up to one month at 4° C in PBS before delipidation.

For CUBIC tissue clearing,^45^ both the spinal cords and brains were delipidated with CUBIC-L consisting of 10 % (wt/wt) N-butyldiethanolamine (Thermo Fisher Scientific, L09953.36) and 10 % (wt/wt) Triton X-100 (Millipore Sigma, 648466) in ddH_2_O. The tissue was immersed in 10 mL CUBIC-L at 37° C overnight with gentle shaking. Then, the tissue was incubated twice similarly for two days each in CUBIC-L. The tissue was washed four times for 2 hours with 10 mL room-temperature PBS to remove surfactants.

The refractive index of tissues was matched to the immersion liquid using CUBIC-R consisting of 45 % (wt/wt) Antipyrine (Thermo Fisher Scientific, 104975000) and 30 % (wt/wt) Nicotinamide (Thermo Fisher Scientific, A15970.30) in ddH_2_O. In experiments involving the detection of native mEmerald or sfGFP, 250 µL N-Butyldiethanolamine (Thermo Fisher Scientific, L09953.36) was added per 50 mL of CUBIC-R to modify the pH of the solution to avoid quenching these EGFP-derived fluorophores. First, the tissue was immersed in 5 mL of ½-diluted CUBIC-R overnight at room temperature with gentle shaking. The tissue was then incubated twice overnight in CUBIC-R.

The tissue was allowed to sit at room temperature for up to one week before being thoroughly washed in PBS for medium-term storage at 4° C or three 2-hour washes and long-term storage at -20° C in a cryoprotectant solution consisting of 50% (v/v) ethylene glycol (Sigma Aldrich, 324558), 20% (v/v) glycerol (Sigma Aldrich, G5516), 6 % (v/v) 10x PBS, and 24% (v/v) ddH_2_O

CUBIC tissue clearing induces some swelling, so coordinates might not precisely match absolute bregma coordinates in living animals.

#### Light-sheet Imaging and Processing

Samples were imaged on the Cleared Tissue LightSheet (CTLS) system (Intelligent Imaging Innovations). The samples were immersed in a mixture of silicon oil (Gelest, PDM-7040, RI = 1.556) and mineral oil (Sigma-Aldrich, M8410, RI = 1.467) at a volume ratio of 0.674:0.326 for a resultant RI of 1.525. The samples were imaged using the green (488 nm excitation, 525/39 nm emission), red (561 nm excitation, 600/52 nm emission), or far-red (640 nm excitation, 679/41 nm emission) laser and filter combinations as situationally appropriate with objective lens on the system (Zeiss, PlanNeoFluar Z 1×/0.25NA FWD=56 mm) at 5x zoom magnification. With this configuration, the X-Y pixel distance was 1.3 μm. The z step was 3 μm, except when coarsely imaging injection sites, in which case it was 10 μm. The two illumination objectives 5×/0.14NA were used to illuminate the sample from the left and right side of the tissue. Samples were imaged in “low” resolution mode, which does not move the focal point of the illumination. All hardware was controlled by Slidebook software (version 2024.1, Intelligent Imaging Innovations, RRID:SCR_014300), and data was stored on a PowerEdge R740XD Server (Dell) equipped with twelve 16 TB HDDs with RAID configuration.

The output of the microscope was z-stacks in Numpy array binary file format. NPY2BDV (version v.1.0.8, DOI:10.5281/zenodo.6148906) was used to convert the files to hierarchical data format (.h5) with associated metadata (.xml). From here, the BigStitcher^136^ plugin (version 1.2.10; https://github.com/PreibischLab/BigStitcher) in Fiji^137^ (RRID:SCR_002285) was used to select left/right illumination tiles, stitch tiles, make minor adjustments in the orientation of the files to the desired plane, and correct for chromatic aberration. After exporting images as .tiff files, they were trimmed to uniform dimensions, their contrast was equalized, and they were combined into one z-stack for the remainder of the analysis in the case of tdT imaging or into separate files in the case of retrograde tracing experiments, as the latter differed more substantially in background fluorescence.

#### Image Segmentation and Object Counting

Image segmentation was performed with Labkit^138^ (release 0.3.11, https://github.com/juglab/labkit-ui) computer vision system with GPU acceleration. We created three labels corresponding to neurons, tissue autofluorescence, and imaging medium to train the pixel classification neural network. We kept adding annotation masks until the prediction satisfied a thorough side-by-side comparison with the original images, erring on the side of conservative annotation to avoid false positives. Additionally, five pairs or triplets of adjacent neurons were annotated per sample to ensure the pixel classifier avoided merging adjacent neurons. With the resultant probability maps, a 3D Gaussian blur (σ of 1:1:1) was used to eliminate stray pixels that would increase the computational burden. From here, the 3DObjectsCounter plugin^139^ (version 2.0.1, https://imagej.net/plugins/3d-objects-counter) in Fiji was used to assign coordinates to the putative neurons with an intensity cutoff chosen using sideby-side comparisons with the original images.

### Anterograde Monosynaptic Tracing

#### Viral Injections

General surgical procedures were followed, as above, with the following specifications: the surgeries were performed on 73-day-old heterozygous Ai14^42^ (The Jackson Laboratory, #007914; RRID:IMSR_JAX:007914). 600 nL of pENN.AAV.hSyn.Cre.WPRE.hGH packaged in AAV1 (Addgene 105553-AAV1, RRID:Addgene_105553) at 1.9e13 vg/mL titer injected at a rate of 100 nL/minute into the caudal forelimb area (CFA) according to the previously-published^86^ coordinates expressed relative to bregma as [mm anterior-posterior, mm medial-lateral, mm dorsal-ventral]: [0.0, 1, -0.65], [0.5, 1, -0.65], [0.5, 1.5, -0.65]. Mice were allowed to recover and express the virus for five weeks before euthanasia. The tissues were cleared imaged on a light-sheet microscope according to the abovementioned procedures.

#### Light-sheet Imaging Data Analysis

General CUBIC tissue clearing and light-sheet imaging procedures were followed, as above, with the following specifications: to align the counted objects of the naturally-curved spinal cord into rectilinear and directly comparable coordinates, virtual “slices” every 25 coronal pixels (130 μm) were subsampled from the data, and the three-dimensional coordinates of the central canal were manually annotated for each slice, as were the length and width of the grey matter. Each point was centered to its nearest annotated midline point, and coordinates were normalized to the length and width of their nearest spinal cord slice grey matter. tdTomato-positive cells were counted as described above. We annotated the data conservatively, yielding a false positive rate against manual annotation of 0% (0/212). This resulted in a false negative rate against manual annotation of 11.7 ± 1.3%. Any counted objects centered less than 10 pixels (52 μm) away from the midline were excluded from the analysis to prevent accidental misassignment of abundant contralateral post-synaptic neurons to the ipsilateral side.

Ridgeline plots were generated with the stat_density_ridges function of the ggridges package (version 0.5.5, DOI: 10.32614/CRAN.package.ggridges). Paired two-sided t-tests were performed using the mean dorso-ventral neuron coordinate of each biological replicate (mouse). No significant sex differences were detected.

### BDA Anterograde Tracing

#### BDA Injections and Histology

Sample injection and histology of mice 1120D (4D), 1120E (5E), and 1120F (6F) occurred in a previous study, which describes the injection and histological procedures.^46^ Briefly, wild-type Balb/C mice (The Jackson Laboratory, #000651; RRID:IMSR_JAX:000651) >90 days of age were injected with 10% (w/v) unconjugated BDA (MW: 10,000, Invitrogen D1956) dissolved in 0.9% sterile saline over 3–4 minutes at the following coordinates expressed relative to bregma as [mm anterior-posterior, mm medial-lateral, mm dorsal-ventral]: [0.0, 1, -0.5]. Mice were allowed to survive for 14 days after surgery, perfused with 4% PFA in 0.1M phosphate buffer, and stored in 4% PFA before being transferred to 27% sucrose at 4° C for one day for cryoprotection. Tissue was embedded in TissueTek ® O.C.T. (Sakura, 4583), and 30 μm sections were taken every 500 μm throughout the length of the spinal cord.

For visualization of BDA staining, sections were washed in PBS and incubated in 1:200 streptavidin-HRP (Perkin Elmer NEL75000-1ea) in PBS for two hours, washed three times in PBS. Then, tyramide signal amplification was performed in 1:100 TSA-Cy3 in the supplied commercial reaction buffer (Perkin Elmer SAT704A001ea). Sections were washed twice in PBS, mounted on gelatin-slubbed slides, and cover slipped with Vectashield mounting medium (Vector Labs H-1000).

#### Imaging and Image Analysis

Spinal cord sections were imaged on an IX80 epifluorescence microscope (Olympus) using the Cy3 filter (545/30 nm excitation, 620/60 nm emission) using a 10x objective. Images were first aligned to their central canals in Fiji (RRID:SCR_002285) and cropped to the length and width of the grey matter of each section. Their dimensions were standardized by cropping and aligning the spinal cords in Krita (version 5.1.5, https://krita.org/) and exporting them as equivalent pixel dimensions before binarization with Fiji. The images were then converted to matrices of binarized pixels with a custom Python script (version 3.13, RRID:SCR_006903). To make the data comparable along the length of the spinal cord, the units were converted to relative distances based on the length and width of the horns. Only pixels lateral to the dorsal funiculus were counted to avoid tabulating the tracts and prevent erroneous mis-assignment of pixel laterality.

Pixel densities and dorso-ventral pixel distributions were plotted with the geom_raster function of the ggplot2 package (version 3.4.4; RRID:SCR_019186). Ridgeline plots were generated with the stat_density_ridges function of the ggridges package (version 0.5.5, DOI: 10.32614/CRAN.package.ggridges). Paired two-sided t-tests were performed using the mean pixel dorso-ventral coordinate of each biological replicate (mouse). No significant sex differences were detected.

### Single-Nucleus Sequencing

#### Viral Injections

General surgical procedures were followed, as above, with the following specifications: the surgeries were performed on 55–71-day-old heterozygous INTACT mice^48^ (The Jackson Laboratory, #021039; RRID:IMSR_JAX:021039). 600 nL of pENN.AAV.hSyn.Cre.WPRE.hGH packaged in AAV1 (Addgene 105553-AAV1; RRID:Addgene_105553) at 1.9e13 vg/mL titer injected at a rate of 100 nL/minute into the caudal forelimb area (CFA) according to the previously-published^86^ coordinates expressed relative to bregma as [mm anterior-posterior, mm medial-lateral, mm dorsal-ventral]: [0.0, 1, -0.65], [0.5, 1, -0.65], [0.5, 1.5, -0.65]. Mice recovered and expressed the virus for six to eight weeks before euthanasia.

#### Sample Processing and Nuclei Isolation

Up to reverse transcription steps, all work was completed with precautions against exogenous RNAses: equipment was cleaned with RNaseZap™ RNase Decontamination Solution (Invitrogen, AM9780) and rinsed with UltraPure™ water (Invitrogen, 10977015), all reagents were certified RNAse-free unless otherwise noted, and work completed on ice.

Mice were deeply anesthetized with a ketamine and xylazine mixture in normal saline (100 mg/kg and 20 mg/kg, respectively). They were then perfused with an artificial cerebrospinal fluid-based buffer comprising 110 mM NaCl (Invitrogen, AM9760G), 8.4 mM HEPES buffer (Thermo Fisher Scientific, 15630080), 25 mM glucose (Sigma Aldrich, G7528-250G), 75 mM sucrose (Fisher Scientific, AC419762500), 8 mM MgCl^2^ (Thermo Fisher Scientific, AM9530G), and 2.5 mM KCl (Thermo Fisher Scientific, AM9640G) (pH = 7.4). The solution was carbogenated on ice for 45 minutes before perfusion. The cervical spinal cord was removed and delaminated, rinsed thoroughly in carbogenated ACSF, and flash-frozen above liquid nitrogen vapor. The time from perfusion to flash freezing was <10 minutes for all samples. Samples were stored at -80° C in tubes with Tissue-Tek® OCT solution (Sakura, 4583) at the bottom of the conical tube to prevent the samples’ desiccation.

To separate ipsilateral from contralateral sides of the spinal cord, the samples were warmed to -20° C overnight and placed in a spinal cord cutting block (AgnThos, 69-4170) also at -20° C. Room temperature razor blades were used to slice the cord into coronal “discs” of 1mm thickness. Again, using the spinal cord matrix to align the razor blades, the spinal cord was cut into three sections under a dissection microscope: ipsilateral, medial, and contralateral. The medial section was thin, existing only to separate the portions of the spinal cord that were clearly ipsilateral from those clearly contralateral. The grey matter near the medial section of each coronal hemi-disk was examined to ensure that the cut was consistent.

To purify nuclei, we used a modified density gradient centrifugation protocol similar to those previously published.^140–142^ “Buffer A” consisted of OptiPrep™ 60% iodixanol (Sigma-Aldrich, D1556). “Buffer B” consisted of 900 mM KCl (Invitrogen, AM9640G), 30 mM MgCl_2_ (Invitrogen, AM9530G), 120 mM Tricine-KOH (pH = 7.8) (Thermo Scientific, 15426389) in UltraPure™ water (Invitrogen, 10977015). “Buffer C” consisted of 5 volumes of Buffer A, 1 volume of Buffer B, 0.5 mM spermidine (Thermo Scientific, A19096.03), 0.15 mM spermine (Sigma-Aldrich, S3256), 1 mM DTT (Thermo Scientific, R0861), and 20 units/mL SUPERase•In™ RNase inhibitor (Invitrogen, AM2694), and 1 tablet per 10 mL of cOmplete™ EDTA-free Protease Inhibitor Cocktail (Roche, 11836170001). “Buffer D” consisted of 0.25 M sucrose (Thermo Scientific, AC419762500), 150 mM KCl (Invitrogen, AM9640G), 5 mM MgCl_2_ (Invitrogen, AM9530G), 20 mM Tricine-KOH (pH 7.8) (Thermo Scientific, 15426389), 0.5 mM spermidine (Thermo Scientific, A19096.03), 0.15 mM spermine (Sigma-Aldrich, S3256), 1 mM DTT (Thermo Scientific, R0861), 20 units/mL SUPERase•In™ RNase inhibitor (Invitrogen, AM2694), and 40 units/mL RNAsin Plus RNase inhibitor (Promega, N2611). “Buffer E” consisted of 1.17 parts Buffer C to 1 part Buffer D. In each buffer, DTT, spermine, and spermidine were added immediately before commencing purification. Buffer E was also mixed immediately before purification.

Tissue pieces for each replicate were added to 4.5 mL of Buffer D in a dounce homogenizer (DWK Life Sciences, 357542) at 4° C, dounced 40 times on ice with the loose (0.089 - 0.14 mm clearance range), and then 40 times with the tight (0.025 - 0.076 mm clearance range) pestles, taking care not to cause bubbles. The result was placed in a 15 mL protein LoBind® conical tube (Eppendorf, 0030122216). The total liquid was brought to 5 mL, and 4.6 mL Buffer C was added. The mixture was inverted ten times. 2 mL of Buffer E was added to a 13.2 mL thin-wall clear centrifuge tube (Beckman Coulter Life Sciences, 344059), and the 9.6 mL of solution was carefully layered on top 1 mL at a time. The solution was spun at 10,000 rpm (max acceleration, level 7 deceleration, max 17,136 g, average 12,320 g, K-factor 2,089) at 4°C in a swinging bucket rotor (SW41) for 30 minutes in a Beckman Coulter XL-70 ultracentrifuge. The supernatant was emptied from the tube by inverting it and “flicking” the liquid out. The nuclei were resuspended in 1 mL of Buffer D with 1:10,000 DAPI and passed through a cell strainer (Corning Life Sciences, 352235) into a 1.5 mL protein LoBind® tube (Eppendorf, 022431064) before being quantified on a Countess II FL Cell Counter (Thermo Fisher Scientific).

#### Fluorescence-Activated Cell Sorting (FACS)

96-well twin.tec® DNA LoBind plates (Eppendorf, 0030129504) of Smart-seq3^50^ lysis buffer were prepared in advance. Lysis buffer consisted of 5% poly-ethylene glycol 8000 (PEG) (Sigma Aldrich, 89510-250G-F), 0.1% Triton X-100 (Sigma Aldrich T8787-50ML), 0.5 µM OligodT30VN (HPLC purification, IDT, /5Biosg/ACGAGCATCAGCAGCATACGAT-TTTTTTTTTTTTTTTTTTTTTTTTTTTTTVN, where /5Biosg/ indicates 5’ Biotin modification, V indicates A, C, or G, and N indicates A, C, G, or T), 0.5 mM each of dNTPs (Thermo Fisher Scientific, R0182), and 0.5 U/µL RNAse Inhibitor (Takara Bio, 2313A) in UltraPure™ water (Invitrogen, 10977015). 3 µL of lysis buffer was deposited via repeater pipette, spun down, covered in a Nunc™ foil seal (Thermo Fisher Scientific, 232698), and stored at -80° C.

Nuclei were sorted on an MA900 Cell Sorter (Sony) with a 130 µm chip (LE-C3213), as the larger droplets facilitated accurate targeting to each well. Before each sort, the machine’s droplet placement was calibrated to ensure accurate droplet placement. The first DAPI peak was gated for each sample to exclude all non-singlet nuclei. Then, using a littermate control, gates were established on the sfGFP signal (channel FL1, 488 nm excitation, 525/50 nm emission) such that no nuclei in a sample of 10,000 were positive. All nuclei above this signal were considered sfGFP positive in subsequent sorts. Nuclei were sorted into plates on a chilled metal base to prevent RNA degradation, then immediately sealed with Nunc™ foil seals (Thermo Fisher Scientific, 232698), spun down 1000 g for 4 minutes at 4° C in a swinging bucket centrifuge, and flash frozen on liquid nitrogen. Samples were stored at -80° C until lysis.

For images of sorted nuclei, sfGFP(+) and sfGFP(-) nuclei were spun onto a slide using a Cytospin™ 4 Cytocentrifuge (Epredia™) and coverslipped using ProLong™ Glass Antifade Mountant (Thermo Fisher Scientific, P36980). They were imaged on an LSM700 Laser Scanning Microscope System (Zeiss) using the pre-configured DAPI and EGFP laser and filter combination on the ZEN Black Edition microscopy software (version 8.1.9.484, RRID:SCR_018163) with a 63x oil immersion objective (Zeiss, Plan-Apochromat 63x/1.4 Oil DIC M27, FWD=0.19mm). For visualization of representative sorts, data was processed using the ggCyto package^143^ (version 3.19, DOI: 10.18129/B9.bioc.ggcyto) and density heatmaps visualized using the geom_bin_2d package of the ggplot2 package (version 3.4.4; RRID:SCR_019186).

#### Smart-seq3

Smart-seq3 was performed as previously described (DOI: 10.17504/protocols.io.bcq4ivyw),^50^ with some changes: plastics and indexing schema were modified for convenience and cost, and we used empirically determined numbers of PCR cycles as the protocol suggests. We did not use 0.2% SDS to strip the Tn5 enzyme after tagmentation. In addition to maintaining an RNAse-free work environment and reagents as previously described, freeze-thaw cycles on all reagents were <2, and all steps were performed at 4° C unless otherwise noted.

Nuclei were lysed in a thermocycler at 72 °C for 10 minutes, with a 4 °C hold. Then, 1 µL of reverse transcription buffer was deposited with a repeat pipetter and spun down for 30 seconds at 4 °C with a tabletop plate centrifuge (LabNet, C1000). The reverse transcription buffer consisted of 25 mM Tris-HCl (pH = 8.3) (Sigma Aldrrich, T6791-100G), 30 mM NaCl (Invitrogen, AM9760G), 2.5 mM MgCl_2_ (Invitrogen, AM9530G) 1 mM GTP (Thermo Fisher Scientific, R1461), 8 mM DTT (Thermo Fisher Scientific, 707265ML), 0.5 U/µL RNAse Inhibitor (Takara Bio, 2313A), 2 µM Template-Switching Oligo (RNAse-free HPLC purification, IDT, /5Biosg/AGAGACAGATTGCGCAATGNNNNNNNNrGrGrG, where /5Biosg/ indicates 5’ biotin modification, r indicates RNA nucleotide, and N indicates A, T, C, or G and is the Unique Molecular Identifier [UMI]), and 2 U/µL Maxima H-minus RT enzyme (Thermo Fisher Scientific, EP0751) in UltraPure™ water (Invitrogen, 10977015). Reverse transcription was performed with a thermocycler at 42 °C for 90 min, followed by ten cycles of 2 minutes at 50 °C and 2 minutes at 42 °C, and enzyme inactivation at 85 °C for 5 minutes. The reaction was then held at 4 °C.

Pre-amplification PCR was performed by depositing 6 µL of reaction mix into each library and spinning down for 30 seconds at 4 °C using a tabletop plate centrifuge (LabNet, C1000). The pre-amplification reaction mix consisted of 1X KAPA HiFi Hotstart buffer (Roche, KK2502), 0.3 mM each of dNTP mix (Thermo Fisher Scientific, R0182), 0.5 mM MgCl_2_ (Thermo Fisher Scientific, AM9530G), 0.5 µM forward primer (HPLC purification, IDT, TCGTCGGCAGCGTCAGATGTGTATAAGAGACAGATTGCGCAA*T*G, where _* indicates a phosphorothioated base), 0.1 µM reverse primer (HPLC purification, IDT, ACGAGCATCAGCAGCATAC*G*A, where _* indicates a phosphorothioated base), and 0.02 U/µL KAPA HiFi DNA Polymerase (Roche, KK2502). The reaction was performed in a thermocycler by denaturing once at 98 °C for 3 minutes, 23 cycles of 98 °C for 20 seconds, 65 °C for 30 seconds, and 72 °C for 4 minutes, and one final elongation incubation at 72 °C for 5 minutes. The reaction was then held at 4 °C. The number of PCR cycles was determined empirically by cycling nuclei 18–25 cycles and examining the resultant cDNA on a high-sensitivity DNA electrophoresis chip (Agilent, 5067-4626). The resultant cDNA was purified with a 0.6:1 ratio of Sera-Mag Speed Beads (Cytiva, 65152105050250) resuspended in 22% PEG as per the mcSCRB-seq protocol^144^ (DOI: 10.17504/protocols.io.p9kdr4w). Briefly, 1 mL of beads were washed twice with and resuspended in a buffer of 10 mM Tris-HCl (pH = 8.0) (Sigma Aldrrich, T6791-100G) and 1mM EDTA (Thermo Fisher Scientific, AM9260G). These beads were resuspended in 50 mL of the PEG suspension buffer, consisting of 22% (w/v) PEG 8000 (Sigma Aldrich, 89510-250G-F) in 1 M NaCl (Invitrogen, AM9760G), 10 mM Tris-HCl (pH = 8.0) (Sigma Aldrrich, T6791-100G), 1 mM EDTA (Thermo Fisher Scientific, AM9260G), 0.01% IGEPAL® (Sigma Aldrich, I8896), and 0.05% (w/v) sodium azide (Sigma Aldrich, S2002-100G). The cDNA was incubated in a 0.6:1 ratio of PEG beads to sample, allowed to incubate at room temperature for 8 minutes, magnetically separated, washed in 100 µL freshly prepared 80% (v/v) ethanol, and again separated. The ethanol mixture was aspirated, and the beads were allowed to dry for 5 minutes before elution.

cDNA was eluted in 12 µL UltraPure™ water, magnetically separated, and the water removed from the beads (Invitrogen, 10977015). cDNA was measured and diluted to 100 pg/µL with the QuantiFluor® dsDNA quantification kit (Promega, E2670) according to the manufacturer’s instructions. cDNA was checked in a randomly chosen well from each plate with a high-sensitivity DNA Bioanalyzer chip kit (Agilent, 5067-4626).

For tagmentation, we first made 4x tagmentation buffer consisting of 40 mM Tris-HCl (pH=7.5) (Sigma Aldrrich, T6791-100G), 20 mM MgCl_2_ (Invitrogen, AM9530G), and 20% (v/v) dimethylformamide (Sigma Aldrich, D4551) in UltraPure™ water (Invitrogen, 10977015). The tagmentation reaction mix consisted of 1X tagmentation buffer and 8% (v/v) Nextera amplicon tagmentation mix (Illumina, FC-131-1096) in UltraPure™ water (Invitrogen, 10977015). 1 µL of 100 pg/µL cDNA was added to 1 µL of tagmentation reaction mix and quickly spun down for 30 seconds at 4 °C using a tabletop plate centrifuge (LabNet, C1000). The reaction was incubated in a thermocycler at 55 °C for 10 minutes. We did not use 0.2% SDS to strip the Tn5 as suggested because the Nextera Tn5 enzyme did not require it. This may have harmed downstream library complexity, but it was done because SDS seemed to inhibit downstream reactions.

Our indexing scheme used combinatorial rather than unique dual indexing to reduce costs. This decision was made based on data suggesting the NextSeq platform, which does not employ ExAmp chemistry, had workably low levels of index hopping.^145^ Our unique dual index pairs were generated with a list of verified barcodes generated in a separate publication.^146^ We added 0.75 µL of each indexing primer to the tagmented cDNA (HPLC purification, IDT, i7: CAAGCAGAAGACGGCATACGAGATnnnnnnnnGTCTCGTGGGCTCGG, i5: AATGATACGGCGACCACCGAGATCTACAC-nnnnnnnnTCGTCGGCAGCGTC, where n indicates a pre-determined index). Then, we added 3 µL of PCR mix to each reaction, consisting of 1X Phusion™ High-Fidelity Buffer (Thermo Scientific, F530L), 0.2 mM each of dNTP mix (Thermo Fisher Scientific, R0182), and 0.01 U/µL o Phusion™ High-Fidelity Enzyme (Thermo Fisher Scientific, F530L) in UltraPure™ water (Invitrogen, 10977015). The reaction proceeded in a thermocycler with a 72 °C incubation for 3 minutes, then a 98 °C incubation for 3 minutes, then 12 cycles of 98 °C for 10 seconds, 55 °C for 30 seconds, and 72 °C for 30 seconds, and a single final incubation of 72 °C for 5 minutes. The reaction was held at 4 °C.

The cDNA was pooled and cleaned up similarly to previous steps using 0.6:1 Sera-Mag Speed Beads (Cytiva, 65152105050250) in 22% PEG. The libraries were eluted in 40 µL UltraPure™ water (Invitrogen, 10977015). Finally, primer-dimers were excluded by running the library on a 2% Agarose E-Gel™ EX (Thermo Fisher Scientific, G402002) and purifying the 550–2kb band with a Qiagen QIAQuick® gel extraction kit (Qiagen, 28704), both per manufacturer’s instructions.

Libraries were sequenced on a NextSeq 500 (Illumina) on two 75-base single-read high-output runs per manufacturer instructions (Illumina, 20024906). As expected, the library fragment size was quite large, so loading concentrations were based on the portion of the fragment distribution likely to efficiently cluster on the NextSeq500 (150–750 bp).^147^ The percent of reads passing the filter was notably low at ∼30% in the first run and ∼46% in the second run. While we are not certain, it appears this was due to the homogeneous sequence before the UMI of each read, which made cluster discrimination inefficient on the NextSeq 500’s flow cell. Reducing loading concentration from 2.7 pM to 1.6 pM and increasing PhiX standard (Illumina, FC-110-3002) concentration from 10% to 30% seemed to help improve cluster discrimination in the second sequencing round.

#### Smart-seq3 Data Processing

We used bcl2fast2 (version 2.20; RRID:SCR_015058) with no lane splitting to convert binary base-call files to fastq. zUMIs^148^ (version 2.9.7; RRID:SCR_016139) was used to process the remainder of the data. We used a barcode “whitelist” given to zUMIs to align only known barcodes. Barcodes were filtered to 3 bases under a phred score of 20 and UMIs to a score of 2 under 20. The hamming distance for binning of barcodes was set to 1, and only barcodes with > 1000 reads were processed. For alignment of reads and processing into error-corrected UMI counts, we created a genome index with STAR^149^ (version 2.7.9a; RRID:SCR_004463) using GEN-CODE’s M27 mouse assembly (https://www.gencodegenes.org/mouse/release_M27.html). Because data was from nuclei, counts from both intronic and exonic reads were used for analysis. From this point, data were processed and mainly visualized in Seurat^150^ (version 5.0.1; RRID:SCR_016341). Individual cells were quality controlled by excluding those with less than 1500 genes or more than 9000 and those with >2% mitochondrial genes. For initial visualization of the data set (Fig S1), nuclei were integrated by their spinal cord side of origin using the SelectIntegrationFeatures (n = 5000) and IntegrateData functions in Seurat. The nuclei were then coarsely clustered (resolution = 0.5) using the FindClusters function. Single-cell heatmaps were generated using the DoHeatmap function of Seurat^150^ (version 5.0.1; RRID:SCR_016341).

#### Spinal Cord Atlas Generation

Mice were anaesthetized with 2.5–3% isoflurane and transcardially perfused with cold, pH 7.4 HEPES buffer containing 110 mM NaCl (Sigma Aldrich, S5886), 10 mM HEPES (Sigma Adlrich, H3375), 25 mM glucose (Sigma Aldrich, G7528), 75 mM sucrose (EMD Millipore, 573113), 7.5 mM MgCl2 (Sigma Aldrich, M1028), and 2.5 mM KCl (Quality Biological, 351-044-101) to remove blood (see DOI: 10.17504/protocols.io.5jyl8peq8g2w/v1). Following perfusion, the spinal cord was dissected quickly, cut into 4 parts (C1-C8, T1-T13, L1-L6, S1-S4 + Co1-Co3), frozen for 2 min in liquid nitrogen vapor and then moved to −80 °C for long term storage following a freezing protocol developed at the Allen Institute for Brain Science (DOI: 10.17504/protocols.io.j8nlkodr6v5r/v1).

Nuclei were isolated using the RAISINs method^151^ with a few modifications as described in a nuclei isolation protocol developed at the Allen Institute for Brain Science (DOI: 10.17504/protocols.io.4r3l22n5pl1y/v1). In short, excised tissue dissectates were transferred to a 12-well plate containing CST extraction buffer and passed through a 0.22 µm filter (Millipore Sigma, SE1M179M6). Mechanical dissociation was performed by chopping the dissectate using spring scissors in ice-cold CST buffer for 10 min. The entire volume of the well was then transferred to a 50-ml conical tube while passing through a 100-µm filter, and the walls of the tube were washed using ST buffer. Next, the suspension was gently transferred to a 15-ml conical tube and centrifuged in a swinging-bucket centrifuge for 5 min at 500 rcf and 4 °C. Following centrifugation, the majority of supernatant was discarded, and pellets were resuspended in 100 µl 0.1x lysis buffer and incubated for 2 min on ice. Following the addition of 1 ml wash buffer, samples were gently filtered using a 20-µm filter (Greiner BioOne, 542120) and centrifuged as before. After centrifugation, most of the supernatant was discarded, pellets were resuspended in 10 µl chilled 1X nuclei buffer (10x Genomics, 2000153) with 1 mM DTT and 1 U/µL RNase inhibitor, and nuclei were counted to determine the concentration. Nuclei were diluted to a concentration targeting 5,000 nuclei per µl.

For 10x Multiome processing, we used the Chromium Next GEM Single Cell Multiome ATAC + Gene Expression Reagent Bundle (1000283, 10x Genomics). We followed the manufacturer’s instructions for transposition, nucleus capture, barcoding, reverse transcription, cDNA amplification, and library construction (DOI: 10.17504/protocols.io.bp2l61mqrvqe/v1). Processing of 10x Genomics snRNA-seq libraries was performed similarly to previous publications.^111,152^ In brief, libraries were sequenced on the Illumina NovaSeq6000, and raw sequencing data were processed using cellranger-arc (10x Genomics; https://www.10xgenomics.com/; version 2.0.2) and the GRCm39 (GCF_000001635.27-RS_2023_04) mouse reference genome to generate single-nucleus RNA-seq (snRNA-seq) UMI count matrices for intronic and exonic reads. Nuclei were filtered to remove lowquality samples by requiring ≥1000 genes detected per non-neuronal nucleus and requiring ≥2000 genes detected per neuronal nucleus. Putative doublets were removed by setting a threshold of 0.3 on doublet score computed by DoubletFinder.

#### Spinal Cord Atlas Integration

We performed integration and clustering analysis among four data sets using Seurat^150^ (version 5.0.1; RRID: SCR_016341). The first was a spinal cord atlas^53^ consisting of data sets from several sources^52–59^ (accessions/identifiers: Russ GSE158380; Sathyamurthy GSE103892; Hayashi GSE108788; Zeisel SRP135960; Häring GSE103840; Rosenberg GSE110823; Baek GSE130312). The second data set profiled ChAT-enriched nuclei (accession: GSE16759).^52^ The functions merge and SplitObject were first used to create lists of Seurat objects for downstream analysis. On each data set, we normalized and found 2000 features using NormalizeData and FindVariableFeatures. Then, we used the function SelectIntegrationFeatures to choose the features to use when integrating data sets. This function ranks features by the number of data sets they are deemed variable in, breaking ties by the median variable feature rank across data sets. It returns the top-scoring features by this ranking. FindIntegrationAnchors was used to find a set of anchors between a list of Seurat objects. These anchors were later used to integrate the objects using the IntegrateData function. Within the integration space (setting DefaultAssay to “integrated”), we conducted clustering (resolution = 0.8), and cells within the same cluster of motor neurons from external data sets are annotated as motor neurons. Finally, we conducted a label transfer from this atlas to the post-synaptic neurons. Neuronal clusters were annotated similarly and confirmed using *Snap25* expression.

#### Data Analysis

First, we limited our analysis to only neuronal populations determined by cluster *Snap25* expression. To determine whether the location of ipsilaterally-tagged post-synaptic neurons in the spinal cord found using our initial trans-synaptic tracing experiment was reflected in our transcriptomic data, we used the cluster location predictions from one of the constituent data sets.^153^ In each cluster, we tabulated the proportion of nuclei from Russ *et al*. 2020’s predicted to be dorsal and mid-to-ventral. We then transferred these labels to our clusters based on which spatial location category was the majority. We manually assigned labels for some neurons with known locations, such as motor neurons, cholinergic interneurons, and CSF-contacting neurons. We similarly tabulated the proportions of putative motor neuronal nuclei and excitatory/inhibitory neuronal nuclei.

Dot plots were generated using the DotPlot function of Seurat^150^ (version 5.0.1; RRID: SCR_016341). Relevant proportions from each biological replicate (pooled or unpooled mice) were subjected to unpaired two-sided t-tests. No significant sex differences were detected, excepting expected sex-associated genes.

### Motor Neuron Monosynaptic Tracing

#### CUBIC Immunohistochemistry

These same samples were used for Figure 1, so methodological information up to the point of immunohistochemical staining is the same. We based our choline acetyltransferase CUBIC immunohistochemistry strategy on a previously published protocol.^73^Following CUBIC-L delipidation and washing, the spinal cord was blocked for two hours at room temperature and gentle rocking in 2.5 mL HEPES-TSC blocking buffer consisting of 5% (v/v) Quadrol (Millipore Sigma, 122262), 10 mM HEPES (Millipore Sigma, 83264), 10% (v/v) Triton X-100 (Millipore Sigma, X100), 200 nM NaCl (Millipore Sigma, S9888), and 0.5% (w/v) casein (Millipore Sigma, C4765) (pH = 7.5). Then, the spinal cord was moved to 2.5 mL HEPES-TSC buffer as above, but with 1:200 dilution of rabbit monoclonal anti-Choline Acetyltransferase antibody conjugated to Alexafluor® 647 (Abcam, ab225262). The spinal cord was incubated with the primary antibodies for five days at room temperature with gentle rocking. Then, it was moved to 4° C for one day on gentle rocking to improve specificity. The sample was then washed twice for one hour each at room temperature in 15 mL of 0.1 M PBT buffer consisting of 85 mM sodium hydrogen phosphate, 15 mM sodium dihydrogen phosphate dihydrate, and 10% (v/v) Triton X-100 (Millipore Sigma, X100) (pH = 7.5). Then, the sample was immersed in 15 mL 1% w/v PFA (Electron Microscopy Sciences, 15700) in 0.1 M PB buffer (as in PBT buffer above, but omitting Triton X-100) (pH = 7.5) for 24 hours on gentle shaking at 25° C. Finally, the sample was washed in 15 ml of 0.1M PB buffer (pH = 7.5) for two hours on gentle shaking at 25° C. Following staining, the refractive index was matched as described above.

#### Light-sheet Imaging and Analysis

General CUBIC tissue clearing and processing procedures were followed, as in Figure 1, with the following specifications: for the annotation of putative motor neurons, the image segmentation was used as a guide, but motor neurons were marked manually by an unblinded experimenter. This was because medial cholinergic interneurons could only be differentiated by their location outside of motor neuron pools, because LabKit had an appreciable false-positive rate in discerning ChAT(+) and tdT(+) cells, and finally because the overall number of tagged neurons was small enough to easily confirm manually. To restrict our analysis to only the regions in immediate proximity to motor neuron pools, we used the two-dimensional kernel density estimation function kde2d from the MASS package (version 7.3-60.0.1, DOI: 10.32614/CRAN.package.MASS) to draw a contour encompassing 95% of all labeled motor neurons, and restricted the data set to this contour. One sample two-sided t-tests (μ = 0.5) were performed on relevant proportions from each biological replicate (mouse). No significant sex differences were detected.

### MAPseq

#### Viral Injections

General surgical procedures were followed, as above, with the following specifications: the surgeries were performed on 289-day-old Ai14^42^ (The Jackson Laboratory, #007914; RRID:IMSR_JAX:007914) heterozygous mice. Two mice included in the analysis were female, and one was male. 300 nL of Sindbis virus library^154^ (MAPseq/BARseq Core in Cold Spring Harbor Laboratory) virus of ∼2e9 vg/mL titer and an estimated 20 million barcode diversity was injected in a square grid composed of the following nine coordinates, expressed relative to bregma as [mm anterior-posterior, mm medial-lateral, mm dorsal-ventral]: [1, 0.5, -0.65], [1.5, 0.5, -0.65], [2, 0.5, -0.65], [2, 0.0, -0.65], [1.5, 0.0, -0.65], [1, 0.0, -0.65], [1, -0.5, -0.65], [1.5, -0.5, -0.65], [2, -0.5, -0.65]. The mice were monitored closely, and the virus was allowed to express for 44 hours before euthanasia.

#### Sample Preparation

Until reverse transcription steps, all work was completed with precautions against exogenous RNAses: equipment was cleaned with RNaseZap™ RNase Decontamination Solution (Invitrogen, AM9780) and rinsed with UltraPure™ water (Invitrogen, 10977015), all reagents were certified RNAse-free, and work completed on ice.

The mice were deeply anesthetized with a ketamine and xylazine mixture in normal saline (100 mg/kg and 20 mg/kg, respectively). The mice were perfused with 50 mL ice-cold carbogenated ACSF as previously, albeit with 1.25% w/v methylene blue (Fisher Scientific, M291-25) to aid in identifying gray matter for spinal cord dissection.^155^ To avoid contamination from endothelial tissue, as observed in the monosynaptic tracing experiment, spinal cords were removed by hydraulic extrusion, which also removes endothelial tissue.^156^ Absent endothelial tissue, the spinal cord easily separates into two sides. Methylene blue-dyed vasculature and gray matter were used to ensure a clean separation of spinal halves. The spinal cord was trimmed into only the cervical segments using the cervical enlargements as a guide. The whole cortex was removed for both the injection site and the opposing cortical hemisphere, and the ipsilateral and contralateral olfactory bulbs were removed as negative controls, and the surface of the tissue carved off. The spinal cord halves were frozen in liquid nitrogen vapor for 1 minute and stored at -80° C. The total time from euthanasia to flash freezing was <10 minutes.

For quality control, the amount of GFP in each sample was quantified with the same reverse transcription reaction described for the library preparations but with EGFP primers (salt purification, IDT, GACGACGGCAACTACAAGAC and TAGTTGTACTCCAGCTTGTGC) and with SYBR Green PowerTrack™ Master Mix (Thermo Fisher Scientific, A46012) according to the manufacturer’s instructions. Samples were compared to a blank negative control and positive controls from past successful sample preparations. Of the eight mice euthanized, one was excluded due to a lost negative control sample, and one was excluded due to a signal in the negative control olfactory bulbs. Four were selected for sequencing based on their high qPCR-inferred barcode expression.

#### Library Preparation and Sequencing

Libraries were prepared as described previously and in the MAPseq protocol (DOI: 10.17504/protocols.io.bsm9nc96):^154^ Tissue was homogenized in 400 μL TRIzol (Thermo Fisher Scientific, 15596026), and then brought to 1 mL total TRIzol. RNA was extracted according to the manufacturer’s protocol and dissolved into 13 μL RNAse-free water.

To begin reverse transcription, 4 μL of dissolved mRNA mixed in 6 μL nuclease-free water, 1 μL spike-in RNA (salt purification, IDT, GTCATGATCATAATACGACTCACTATAGGGGACGAGCTG TACAAGTAAAC-GCGTAATGATACGGCGACCACCGAGATCTACACTCTTTCCCTACACGAC-GCTCTTCCGATCTNNNNNNNNNNNNNNNNNNNNNNNNATCAGTCATCGGAGCGGCCGCTACCTAATTGCCGTCGTGAGGTACGACCACCGCTAGCTGTACA, where ATCAGTCA is the barcode tag of the spike-in and N indicates A, C, G, or T. Spike-in RNA was transcribed in vitro by T7 RNA polymerase and diluted into 10^3^ molecules/ul for target sites and 10^5^ molecules/μL for injection sites) (Integrated DNA Technologies), 1 μL of 10 μM RT primer (salt purification, IDT, CTTGGCACCCGAGAATTC-CANNNNNNNNNNNNXXXXXXXXTGTACAGCTAGCGGTGGTCG, where N indicates A, C, G, or T and is the UMI and X is the sample identifier barcode) (Integrated DNA Technologies), and 1 μL of 10 uM dNTPs. The mixture of RNAs was denatured at 70° C for 10 minutes and cooled on ice for 5 minutes. To complete the reverse transcription, 1 μL SuperScript IV (Thermo Fisher Scientific, 18090010), 4 μL SuperScript IV buffer, 1 μL 0.1 M DTT, and 1 μL RNAsin Plus (Promega, N2611) were added into the mixture, and incubated for 10 minutes at 55°C and 10 minutes at 80°C. Samples were pooled and cleaned with 1.8X AMPure XP beads per the manufacturer’s instructions (Beckman Coulter, A63881).

For second strand synthesis, 17 μL cDNA was added to 2.4 μL SuperScript IV buffer, 0.6 μL 0.1M DTT, 5.6 μL second strand kit buffer (Thermo Fisher Scientific, A48571), 0.75 μL 10mM dNTPs, 0.25 μL *e. coli* DNA ligase, 1 μL DNA polymerase I, and 0.25 μL RNAseH. These were mixed well and incubated for two hours at 16°C, and then 1 μL T4 DNA Polymerase was added and the sample was incubated for 10 minutes again at 16°C. Secondstrand products were cleaned with 1.8X AMPure XP beads (Beckman Coulter, A63881). Following second strand synthesis, 1 μL Exonuclease I (New England Biolabs, M0293S) was added per 16 μL bead-purified product, along with 2 μL of Exonuclease I buffer. The reaction was incubated for 1 hour at 37 °C and inactivated at 80 °C for 20 minutes. All injection sites were pooled together, as were all target sites for the following steps.

The reaction was then PCR amplified by adding 25 μL Accuprime Buffer (Thermo Fisher Scientific, 12346086), 25 μL 10 μM Nested1st gfpF primer (HPLC purified, IDT, CTGTACAAGTAAACGCGTAATG), 25 μL 10 Μm Nested2nd R primer (salt purification, IDT, CAAGCAGAAGACGGCATACGAGATCGTGATGTGACTGGAGTTCCTTGGCACCCGAGAATTCCA), 2.5 μL Accuprime Pfx HF enzyme, and 152.5 μL water. The reaction was incubated at 95ºC for 2 minutes, followed by 15 cycles of 95ºC for 15 seconds and 68ºC for 2.5 minutes. The final extension was at 68ºC for 5 minutes. The samples were again exonuclease digested by adding 5 μL Exonuclease I, incubated at 37 °C for 30 minutes, and inactivated at 80 °C for 20 minutes. The products were diluted 10x for the following PCR steps.

For the final PCR, 2.5 μL of diluted product from the first PCR reaction was added to 2.5 μL Accuprime buffer, 2.5 μL 10 μM Sol I primer (salt purification, IDT, AATGATACGGCGACCACCGA), 2.5 μL 10 μM Sol II primer (salt purification, IDT, CAAGCAGAAGACGGCATACGA), 0.25 μL Accuprime Pfx HF enzyme (Thermo Fisher Scientific, 12346086), and 14.75 μL water. The final PCR cycles were determined by testing each sample in multiples of two cycles and running the resultant product on a gel. The minimal number of cycles to produce a visible and clean band at 230 bp was used. The reaction was incubated at 95ºC for 2 minutes, followed by the empirically determined number of cycles of 95ºC for 15 seconds and 68ºC for 2.5 minutes. The final extension was at 68ºC for 5 minutes.

The product was then cleaned with the Wizard SV Gel and PCR Clean-Up System (Promega, A9282) and eluted into 40 μL per 1 mL of initial pooled PCR product volume. Then, the purified product was loaded onto a 2% agarose gel, and the 230 bp band was cut and purified with a MinElute Gel Extraction Kit (Qiagen, 28604). The 230 bp product was analyzed on a DNA Bioanalyzer Chip (Agilent, 5067-1504) on a Bioanalyzer (Agilent) to confirm the size and concentration for sequencing and then sequenced on a NextSeq 500 (Illumina) on a high-output paired-end 36 bp run (Illumina, 20024906) with the SBS3T sequencing primer (ACACTCTTTCCCTACACGACGCTCTTCCGATCT, Illumina) for paired-end 1 and the Illumina small RNA sequencing primer 2 (CTTGGCACCCGAGAATTCCA, Illumina) for the second paired-end, resulting in 388 million reads passing Illumina standard quality control.

#### Barcode Data Analysis

Raw sequences were first separated according to unique sample identifiers, corresponding to individual mice and dissected brain regions. Data was processed with custom MATLAB (RRID:SCR_001622) scripts. For each brain area with a given barcode and UMI combination, reads were tabulated according to empirically determined thresholds: two for the injection site and five for the target sites. Any barcodes with a hamming distance of no more than three (of thirty bp barcode length) were collapsed. After collapsing reads, data was further filtered to barcodes >50 counts in the injection area and > 5 counts in at least one projection area. Finally, counts were normalized according to the number of spike-in molecules in a given dissected region.

The theoretical estimation of the number of re-used barcodes within a 20 million barcode library is ∼0.1%. The template switching rate induced during sample preparation was estimated to be 0.2%. For analysis, in contrast to previous MAPseq papers, barcode counts were normalized according to the relative number of counts in the injection site. This was because the standard method, in which the barcodes are normalized among several projection sites, was unsuitable due to the data of interest being distributed across only two. One of four samples was excluded from the analysis due to several-fold higher rates of ipsilateral projections, also apparent from the qPCR controls, which was thought to be likely due to contamination during dissection.

Outlier barcodes were discarded using the criteria of 1.5 interquartile ranges away from the third quartile of barcode expression in each dissected anatomical region. This reduced the data to 93.8%–99.3% of its original size, depending on the region. Finally, the data were thresholded based on the negative control olfactory bulb signal. Any normalized barcode value less than the greatest negative control signal was set to 0. One sample, two-sided t-tests (μ = 0.5) were performed on relevant proportions for each biological replicate (mouse). No significant sex differences were observed.

### Retrograde Tracing for Cortical Distribution

#### Viral Injections

General surgical procedures were followed, as above, with the following specifications: vertebrae C4–C5 were removed bilaterally from 99–113 day-old mice to target spinal cord segments C5–C6. At 0.65 mm medial of the midline on the right side, the needle was inserted first 1.1 mm into the cord’s dorsal surface and then withdrawn to 0.9 mm.^157^ 60 nL of AAVrg^7^ (Janelia Viral Tools Core) expressing either mEmerald (RRID:Addgene_54282) or mCherry (RRID:Addgene_55146) driven by a CMV promoter and fused to TOMM20 was injected over 1 minute. The virus was diluted with normal saline to 1.2e13 vg/mL titer. Mice recovered and expressed the virus for three weeks before euthanasia.

#### Tissue Clearing Light-sheet Imaging

Tissue clearing and light-sheet imaging were performed following the general procedures described above.

#### Data Analysis

General CUBIC tissue clearing and processing procedures were followed, as above. Images were annotated as separate samples because viral injections and backgrounds varied between individual replicates. We annotated the data conservatively, yielding a false positive rate against manual annotation of 0.55% ± 0.77% (1/141) in mCherry(+) neurons and 0.0 ± 0.0% (0/160) in mEmerald(+) neurons. This resulted in a false negative rate against manual annotation of 36.0 ± 9.9% (76/216) in mCherry(+) neurons and 38.3 ± 3.0% (100/260) in mEmerald(+) neurons at maximum contrast; however, many of these dim neurons were difficult to discern against background fluorescence and often impossible to discern as separate neurons when adjacent. Double-labeled cells were too infrequent to determine a false positive and negative rate manually, but all were manually confirmed before analysis. The coordinates were centered at approximately the mouse’s Bregma using the Allen Mouse Brain Common Coordinate Framework,^158^ and coordinates were transformed into units of physical distance from the voxel unit of the light-sheet imaging.

One mouse was excluded from the analysis due to subpar imaging quality in S2. All data points were visualized in X-Y, X-Z, and Y-Z coordinates to discern and exclude likely false positives, which were generally the result of autofluorescence at the tissue’s edges. Additionally, extraordinarily large objects relative to the total distribution of volumes (>150 px^3^, >88.76 μm^3^) were excluded as likely false positives. Density plots were generated with the stat_density_2d, and bar plots were generated with the geom_bar functions of ggplot2 (version 3.4.4; RRID:SCR_019186). One sample two-sided t-tests (μ = 0.5) were performed on relevant proportions with Bonferroni corrections for multiple comparisons. No significant sex differences were detected.

### Viral TRAP

#### Viral Injections

General surgical procedures were followed, as above, with the following specifications: vertebrae C4–C5 were removed bilaterally from 164–172-day-old mice to target spinal cord segments C5–C6. At three sites, 0.65 mm medial of the midline on the right side only and 0.5 mm apart rostro-caudally, the needle was inserted first 1.1 mm into the cord’s dorsal surface and then withdrawn to 0.9 mm^157^ The mice were unilaterally injected with AAV2-retro constitutively expressing ribosomal protein L10a fused to EGFP driven by a constitutive Ef1a promoter (Addgene RRID pending) was packaged into AAVrg81,87 (Janelia Viral Vector Core). The virus was diluted in sterile normal saline to 3.45e13 vg/mL titer and injected over 1 minute at each of the three sites. Mice recovered and expressed the virus for 25– 28 days before euthanasia. The cortical distribution of EGFP-L10a(+) neurons was confirmed using the same methods described in previous retrograde tracing light-sheet imaging experiments. In this case, the annotations were not compared to manual counting.

#### Sample Preparation

Until reverse transcription steps, all work was completed with precautions against exogenous RNAses: equipment was cleaned with RNaseZap™ RNase Decontamination Solution (Invitrogen, AM9780) and rinsed with UltraPure™ water (Invitrogen, 10977015), all reagents were RNAse-free, and work completed on ice.

Seven replicates of three pooled mice were processed bilaterally for each region of interest. Both unprecipitated input RNA and polysome immunoprecipitated (IP) samples were prepared for a total of 28 libraries. Two libraries were excluded due to non-Layer 5b and non-neuronal marker expression, presumably due to failed pulldowns. Ultimately, two replicates from male mice and three from female mice were used for the analysis. All 15 spinal cords were cleared with CUBIC tissue clearing and imaged coarsely; however, because the tissue was not perfused with PBS, the autofluorescence from the remaining blood made the injection site less clear than in perfused tissue. An unused mouse was cleared, and the cortex and spinal cord were imaged thoroughly to ensure that the injection protocol produced unilateral expression and the previously observed cortical distribution.

Each mouse was rapidly decapitated, and the cortices ipsilateral and contralateral to the viral injection site were rapidly dissected in an ice-cold buffer consisting of 1X HBSS, 2.5 mM HEPES-KOH (pH = 7.4), 35 mM glucose (Sigma Aldrich, G7528-250G), 4 mM NaHCO_3_ (Sigma Aldrich, S6297-250G), and fresh 100 μg/ml cycloheximide (Sigma Aldrich, C7698-5G) in UltraPure™ water (Invitrogen, 10977015). Each tissue section was rewashed in the dissection buffer to remove excess blood. The triplicate hemicortices were then transferred to tissue homogenizers (DWK Life Sciences, K885510-0020) with 1 mL each of ice-cold homogenization buffer consisting of 10 mM HEPES-KOH (pH 7.4) (Thermo Fisher Scientific, AM9640G), 150 mM KCl (Thermo Fisher Scientific, AM9640G), 5 mM MgCl_2_ (Thermo Fisher Scientific, AM9530G), fresh 0.5 mM DTT (Sigma Aldrich, D9779-5G), 1 tablet per 10 mL cOmplete™ EDTA-free Protease Inhibitor Cocktail (Roche, 11836170001), 400 Units/mL RNasin® Recombinant RNase Inhibitor (Promega, N2511), 200 Units/mL SUPERase•In™ RNase inhibitor (Invitrogen, AM2694), and 100 μg/ml cycloheximide (Sigma, C7698-5G) in UltraPure™ water (Invitrogen, 10977015). Tissues were homogenized with Teflon tissue homogenizing pestles (DWK Life Sciences, K886000-0019, 0.1–0.15mm clearance) using a lab stirrer (Yamato Scientific Co, 231349) for three strokes at 300 RPM. Then, the rotational speed was increased to 900 RPM for 12 additional strokes.

The homogenate was transferred into pre-chilled 1.5 mL centrifuge tubes, spun at 2000 x g for 10 minutes at 4°C, and then transferred to a fresh 1.5 mL centrifuge tube. The homogenate volume was measured at this point, and 1/9th of this volume of 10% NP-40/IGEPAL was added (Sigma, I8896-50ML). Based on the resultant volume of this addition, 1/9th a volume of 300 mM DHPC (Avanti Polar Lipids, 850306P) was added. The samples were gently mixed by inversion five times and incubated on ice for 5 minutes. Then, the samples were spun at 20,000 x g for 15 minutes at 4°C.

#### Ribosomal Pulldown

TRAP was performed as previously,^87^ albeit with a low-yield protocol consisting of half the standard amount of antibodies and beads in anticipation of the potential for differing pulldown efficiencies between neurons of differing abundance. The rationale was that by pooling mice and restricting bead-antibody conjugates, the beads might saturate equally between the differing amounts of available GFP and yield equivalent pulldowns.

The magnetic beads were prepared during the previous day and in tandem with sample preparation. 150 μl for every sample of Streptavidin MyOne T1 Dynabeads (Invitrogen, 65601) were resuspended and washed 1X PBS. Each 500 μg vial of Pierce™ Biotinylated Protein L (Thermo Fisher Scientific, 29997) was resuspended on ice in 500 μl PBS. Each aliquot of beads was resuspended in 880 μl PBS and 120 μl Biotinylated Protein L and allowed to incubate at RT for 35 minutes with gentle end-to-end rotation. The beads were then washed five times with 1 mL 1X PBS containing 3% w/v nuclease-free BSA (Jackson Immunores. 001-000-162) and resuspended in 500 μl wash buffer containing 10 mM HEPES-KOH (pH = 7.4) (Fisher Scientific, AAJ16924AE), 150 mM KCl (Thermo Fisher, AM9640G), 5 mM MgCl_2_ (Thermo Fisher Scientific, AM9530G), 1% (v/v) IGEPAL® (Sigma Aldrich, I8896-50ML), 0.5 mM DTT (Sigma Aldrich, D9779-5G), 40 Units/mL RNasin® Recombinant RNase Inhibitor (Promega, N2511), 20 Units/mL SUPERase•In™ RNase inhibitor (Invitrogen, AM2694), and 100 μg/ml cycloheximide (Sigma Aldrich, C7698-5G) in UltraPure™ water (Invitrogen, 10977015). To this was added 50 μg total of mouse α-EGFP clones 19C8 and 19F7 mAb (25 μg each) (Memorial Sloan-Kettering Antibody and Bioresource Core Facility). The beads were then incubated with slow tilt rotation for >1 hr at RT. During the high-speed spin of the samples, the beads were washed thrice with 1 mL of the aforementioned 0.15 M KCl wash buffer and resuspended in 180 μl. 20 μl 300 mM DHPC (Avanti Polar Lipids, 850306P) was added to the resuspension.

After the high-speed spin, 30 μl of supernatant was removed as “input” samples mixed with 350 μl buffer containing 340 μl RLT and 10 μl βME from the RNeasy Plus Micro kit (Qiagen, 74034) and flash frozen for parallel purification with other samples. The remaining supernatant was placed in a chilled 1.5 mL tube, mixed with 200 μl bead and antibody solution, and incubated on gentle rotation at 4°C overnight.

The following morning, beads were magnetically collected on ice. 30 μl of supernatant was reserved as “unbound” RNA but ultimately not sequenced. The remaining supernatant was removed and washed four times with 1 mL of wash buffer containing 10 mM HEPES-KOH (pH = 7.4) (Fisher Scientific, AAJ16924AE), 350 mM KCl (Thermo Fisher Scientific, AM9640G), 5 mM MgCl_2_ (Thermo Fisher Scienific, AM9530G), 1% (v/v) IGEPAL® (Sigma Aldrich, I8896-50ML), 0.5 mM DTT (Sigma Aldrich, D9779-5G), 40 Units/mL RNasin® Recombinant RNase Inhibitor (Promega, N2511), and 100 μg/ml cycloheximide in UltraPure™ water (Invitrogen, 10977015). All samples, bound and unbound, were then suspended in 350 μl RLT-βME buffer as described above and incubated at room temperature on gentle end-over-end rotation for 10 minutes. The beads were magnetically separated, and the supernatant was removed and placed in fresh 1.5 mL tubes. RNA was purified with the RNeasy Plus Micro kit per the manufacturer’s instructions but eluted into UltraPure™ water (Invitrogen, 10977015) rather than the provided elution buffer (Qiagen, 74034). RNA was quantified, and RNA integrity was confirmed with a Nanodrop spectrophotometer as well as a TapeStation High-Sensitivity RNA ScreenTape (Agilent, 5067-5579).

#### Library Preparation and Sequencing

Libraries were prepared using the Revelo™ RNA-Seq High Sensitivity library preparation kit (Tecan, 30201358) and associated SPIABoost™ Mouse reagents (Tecan, 30201371) and sequencing adapters (Tecan, S02317-FG) per manufacturer’s instructions. Library fragment distribution was analyzed with a TapeStation D1000 DNA ScreenTape (Agilent, 5067-5582). Libraries were sequenced on a NextSeq 500 (Illumina) on a high-output paired-end 75 base pair run (Illumina, 20024907). A median of 35,208,430 paired-end reads passed the filter, ranging from 5,524,727 to 77,421,692 reads per sample.

#### Data Analysis

Binary base-call files were converted to Fastq with bcl2fast2 (version 2.20, RRID:SCR_015058). Sequences were aligned and converted to feature count matrices using a pipeline from Rockefeller University’s Bioinformatics Resource Center (https://github.com/RockefellerUniversity/RU_RNAseq.git).

Sequence and transcript coordinates for mouse mm10 UCSC genome and gene models were retrieved from the BSgenome.Mmusculus.UCSC.mm10 Bioconductor package (version 1.4.0; DOI: 10.18129/B9.bioc.BSgenome.Mmusculus.UCSC.mm10) and TxDb.Mmusculus.UCSC.mm10.knownGene (version 3.4.0; DOI: 10.18129/B9.bioc.TxDb.Mmusculus.UCSC.mm10.knownGene) Bioconductor libraries respectively. Reads were aligned to the genome with Rsubread’s subjunc method^159^ (version 1.30.6; RRID:SCR_016945) and exported as bigWigs normalized to reads per million with the rtracklayer package (version 1.40.6; RRID:SCR_021325; 10.18129/B9.bioc.rtracklayer). Reads in genes were counted with the featurecounts function within the Rsubread package against the full gene bodies (Genebody.Counts) and gene exons (Gene.Counts). This resulted in a median of 87.1% mapped reads ranging from 80.3 to 91.4% per sample.

Normalization, visualization of principal components across experimental groups, and differential expression of counts were performed using DESeq2^160^ (version 1.42.0; RRID:SCR_015687) under default analysis parameters for paired replicates (three pooled mice each). No significant sex differences were observed, excepting expected sex-associated genes. Two replicates were excluded due to the expression of non-neuronal markers, suggesting unsuccessful pulldowns. Paired two-sided t-tests were performed on TPM counts for Rpl10a from each biological replicate (three pooled mice), and this feature was then removed from downstream analysis. To test whether the sets of canonical Layer 5b markers and the sets of Gprin3 bacTRAP-derived Layer 5b markers were equivalently enriched in both pulldowns, we used the CAMERA gene set test procedure^161^ as implemented in the Limma^162^ package (version 3.58.1; RRID:SCR_010943).

Deposited fold-changes and p-values (accession: GSE126957) were used to compare our TRAP expression data to TRAP expression data from injured corticospinal neurons and injured corticospinal neurons treated with neural precursor cells.^95^ To compare our TRAP data to microarray data of developing corticospinal neurons (accession: GSE2039)^99^ deposited microarray data were read with the Affy package^163^ (version 1.80.0; RRID:SCR_012835) and the gcrma package (version 2.74.0; DOI: 10.18129/B9.bioc.gcrma) was used to process optical intensities into expression values. Affymetrix Mouse Expression Set 430 annotation data (version 3.13.0; DOI: 10.18129/B9.bioc.moe430a.db) was used to annotate the probe values. Redundant probes were averaged using Limma’s^162^ (version 3.58.1; RRID:SCR_010943) avereps function.

Rank-Rank Hypergeometric Overlap was performed with the RRHO package^97^ (version 1.42.0; RRID:SCR_014024), but visualizations were produced with the RRHO2^98^ (version 1, RRID:SCR_022754) package, in both instances using default parameters and the Benjamini-Yekutieli correction for multiple comparisons. Genes from each differential expression analysis were ranked using log10(FDR) multiplied by the sign of their fold change. RRHO trims the data to matching genes and removes NAs from the data set, resulting in similar but differing sets of genes compared in each analysis. For this reason, resultant heat map quadrants are not perfectly square. The RRHO package generates lists of concordant genes between two differential expression analyses used for subsequent gene ontology analysis.

Over-representation analysis of gene lists in a gene ontology of biological processes was performed with the clusterProfiler^164^ (version 4.10.0; RRID:SCR_016884) package with a p-value cutoff of 0.01 and Benjamini-Hochberg correction for multiple comparisons. For lists of shared upregulation between two experiments, we used the output files generated for this purpose by the RRHO package. For the comparison of ipsilateral and contralateral IP samples, we used a cutoff of FDR < 0.05. GO results were visualized using enrichPlot (version 1.22.0, DOI: 10.18129/B9.bioc.enrichplot). Venn diagrams were generated using the VennDiagram package (version 1.7.3, DOI: 10.32614/CRAN.package.VennDiagram).

### Retrograde Tracing with Dorsal Hemisection

#### Viral Injections

General surgical procedures as described above were followed, with the following specifications: rAAV2-retro-CAGIG-H2B-V5-mGreenLantern^82^ (Addgene RRID pending, Viral Vector Core, Miami Project to Cure Paralysis) and rAAV2-retro-CAGIG-H2B-V5-mScarlet^82^ (RRID:Addgene_191093, Viral Vector Core, Miami Project to Cure Paralysis), both driven by a CAGIG promoter and of 1.37e14 GC/mL and 2.85e14 GC/mL titer, respectively, were injected into the spinal cords of 48–66 day-old mice on opposite sides of the L2–L4 segments (under vertebrae T12–13) at three sites separated by 0.5 mm rostro-caudally. The segments were located using the apex of the natural curvature of the mouse’s spine and the thickness and spine shape of each vertebra. The lumbar segment of interest was fastened with two custom metal braces inserted into the groove of the spinal vertebrae at this level, and the thick vertebrae were thinned with a rotary tool (Dremel) before being removed with blunt forceps. At 0.65 mm medial of the midline on the right side, the needle was inserted first 1.1 mm into the cord’s dorsal surface and then withdrawn to 0.9 mm.^157^ 60 nL of either virus was injected at approximately 100 nL/minute. Mice were allowed to recover for ten days before the hemisection procedure. Mice demonstrating any hindlimb deficits at this point were excluded and euthanized.

#### Dorsal Hemisections and Sham Surgeries

Mice were randomly assigned hemisection and sham groups, keeping proportions equal within sex and cage. Sample size estimates were based on prior studies, aiming for 10–15 mice per group. The spinal cord was delaminated at the T8 section (vertebra T6), located by counting back from the T2 spinal landmark. While this was fairly consistent, the amount of musculature surrounding the thoracic cord made precise identification of the segment more difficult than in other experiments described. Sham surgeries stopped at this point, whereas the hemisected mice continued.

The hemisection was performed first with very fine scissors (Fine Science Tools, 15000-03) marked at 0.8 mm to make an initial cut through the dura and dorsal half of the spinal cord. To ensure that the spinal cord was hemisected completely, a diamond micro knife (Fine Science Tools, 10100-30) was affixed to a stereotactic frame and used to cut from one side of the spinal cord to the other at a depth of 0.8 mm, three times in either direction of cut. To test the efficacy of the DHX procedure, pENN.AAV.hSyn.Cre.WPRE.hGH packaged in AAV1 (Addgene 105553-AAV1, RRID:Addgene_105553) at 1.9e13 vg/mL titer as injected into a 60-day-old heterozygous Ai14 tdT mouse at a rate of 100 nL/minute into the hindlimb cortex according to the previously-published^46,103^ coordinates expressed relative to bregma as [mm anterior-posterior, mm medial-lateral, mm dorsal-ventral]: [-1.3, 1, -0.65]. After two weeks of recovery, the animal was hemisected as above and allowed to recover for two more weeks before being euthanized, and its spinal cord was cleared and imaged according to general CUBIC tissue clearing and light-sheet imaging procedures described above. In this way, complete transection was visible using tdT fluorescence of the CST.

At this point, hemisected mice display near-complete hindlimb paralysis. Stepping ability recovers hours to days after surgery; however, any mice with hindlimb stepping ability immediately after the surgery were excluded as probable incomplete hemisections. Mice were closely monitored after this surgery, as detailed in general spinal cord surgery protocols, and paper towels were given underneath their bedding to ensure they could easily navigate their cage. Mice that did not resume nest building after several days or mice that lost body weight through recovery were excluded and euthanized, as were mice that failed to regain stepping ability. The mice were allowed to recover for 28 days before fluorogold injections.

#### Fluorogold Injections

Injections of 2% Fluor-Gold^165^ (hydroxystilbamidine)^166^ in sterile normal saline were performed bilaterally at three sites from C3-C4, identified using the beginning of musculature attaching the spinal vertebrae to the mouse’s skull. In contrast to our other cervical injections, we injected 0.5 mm lateral to the midline on either side by moving the needle down 0.9 mm and then retracting it to 0.6 mm. This difference from previous injection sites in the cervical spinal cord was to more closely mimic the methods of the publication on which we were modeling the experiment, adapted to the scale of mice rather than rats by comparing spinal cord atlases of the relevant regions.^106^ 50 nL of 2% Fluoro-Gold was injected at approximately 100 nl/minute.

Fluoro-Gold’s neurotoxic properties at injection sites are well-known.^167,168^ Furthermore, pharmacological investigation suggests that Fluoro-Gold enters neurons via AMPA receptor-mediated internalization.^169^ Others have suggested that Fluoro-Gold enters neurons via endocytosis mediated by a favorable pH gradient irrespective of AMPA receptors.^166^ From our observations, it seems likely that Fluoro-Gold transiently activates motor neurons, directly or indirectly, resulting in rhythmic limb movements for several minutes after injection.

Any mouse experiencing forelimb deficits following this surgery was excluded, as was any mouse that did not resume normal nesting behavior during recovery. Mice were allowed to recover for seven days before being euthanized.

#### Tissue Collection and Processing

Mice were euthanized with a ketamine and xylazine mixture in normal saline (200 mg/kg and 20 mg/kg, respectively). Immediately, the mice were perfused with 30 mL ice-cold PBS, followed by 30 mL ice-cold PBS with 4% paraformaldehyde. The brain and spinal cord were removed and placed overnight at 4° C in 4% paraformaldehyde before being washed three times for two hours in room-temperature PBS. The tissue was then cryoprotected by incubation in 30% sucrose in 1X PBS until the tissue no longer floated in the dense solution.

Brains were frozen in ethanol and dry ice slurry in Tissue-Tek® OCT solution (Sakura, 4583). 50 μM slices were taken from the end of the cerebellum to the rostral-most part of the brain, mounting every third slice after washing in PBS. The slices were allowed to dry and coverslipped with ProLong™ Glass Antifade Mountant (Thermo Fisher Scientific, P36980). The slides were dried at room temperature for 72 hours before being moved to storage at 4° C. One mouse was excluded for insufficient fluorogold signal, two were excluded due to midline-crossing AAV injections, and five sham mice were excluded for gross hindlimb dysfunction from the initial lumbar injection.

#### Imaging

Slices were imaged on an Axio Imager 2 epifluorescence microscope (Zeiss) illuminated with a mercury arc lamp, which provided the ultra-violet spectrum needed for illuminating Fluoro-Gold staining. Additionally, because traditional DAPI filter sets do not capture the wide-band emission spectrum of Fluoro-Gold, we used a custom filter set (350/50 excitation,

>440nm long pass emission) (Chroma, 49025) alongside the microscope’s standard filter sets for mGreenLantern (470/40 nm excitation, 525/50 nm emission) and mScarlet (545/30 nm excitation, 620/60 nm emission) visualization. Images were acquired using a 10x objective (Zeiss, EC PlanNeofluar 10x/0.3 M27, FWD=5.2mm) with a 4000 ms exposure for FluoroGold and a 1000 ms exposure for the two nuclear fluorophores. At this magnification, the distance between pixels was 0.74 μm. A color camera (QImaging, RET-2000R-F-CLR-12-C) in Neurolucida software (version 2021, RRID:SCR_001775) was used to capture the images.

#### Image Processing

For each slice, the red pixel values of the Fluoro-Gold image were used, as they had the lowest background due to tissue autofluorescence. Channels were merged in Fiji (RRID:SCR_002285), and all slices were placed in a stack for image processing and analysis. First, background was subtracted using the rolling ball algorithm (radius of 50 pixels, 37 μm) in Fiji, followed by Gaussian blur (σ = 2) for denoising. There was non-negligible bleed-through from mScarlet into the Fluoro-Gold channel, so we performed spectral unmixing by mouse with the Spectral Unmixing plugin (version 1.20, https://imagej.net/ij/plugins/spectral-unmixing.html). This removed most of the aberrant signal but remained an important potential source of false positives for the remainder of the analysis.

#### Data Analysis

We added the mScarlet and mGreenLantern channels in Fiji to identify neuronal nuclei. We thresholded the image conservatively to a pixel value of 147 out of 255 using a visual comparison to the original images across all samples. The resultant binarized image was separated into distinct non-overlapping nuclei using the watershed algorithm in Fiji with default parameters. Finally, the particles were analyzed with a circularity cutoff of 0.8. We extracted the pixel values in each channel from each labeled nucleus — Fluoro-Gold, mScarlet, and mGreenLantern. We manually found the midline of each slice and recorded it in the metadata file for later centering of the slices. For each mouse, we manually found the coronal slice closest to -1.06 relative to bregma and used it to tabulate the anatomical locations of the serial sections. Some slices were then coordinate-inverted such that the predominant fluorophore in each slice was on the same side of each image. To find the number of total FG(+) neurons irrespective of the labeled nuclei, we used the same annotation procedure, albeit with a cutoff of mean pixel value 55 of 255 and a minimum circularity of 0.5. The circularity threshold for FG(+) neurons was lower than that of nuclei because their cell bodies were less circular than neuronal nuclei.

Abnormally large or small particles (pixel area > 350 pixels^2^, 639.2 μm^2^ or pixel area < 20 pixels^2^, 36.5 μm^2^) were excluded as probable false positives, as were particles far outside the hindlimb cortex. A mean FG pixel value of >55 of 255 was chosen because, on blinded visual examination, it was clearly above the risk of false positives but not yet so strict as to push the numbers of FG(+) neurons 0 in multiple samples. To identify ipsilateral neurons, we manually counted the number of ipsilateral nuclei in 4 mice (2 sham and 2 hemisected) to find 13.5 ± 0.8% ipsilateral. Ipsilateral nuclei were then annotated as the 13.5% of nuclei most expressing their ipsilateral fluorophore in each mouse. To determine the number of FG(+) neurons total in the hindlimb field, we used the two-dimensional kernel density estimation function kde2d from the MASS package (version 7.3-60.0.1, DOI: 10.32614/CRAN.package.MASS) to draw a contour encompassing 95% of all labeled nuclei, and counted the total number of FG(+) neurons within that contour. Fluorogold totals were then calculated for ipsilateral and contralateral neurons and expressed as a proportion of total ipsilateral and contralateral nuclei in each mouse and hemisphere. To compensate for varying FG injection efficiency, these proportions were normalized to the total number of FG(+) neurons in the hindlimb field in its hemisphere. The two hemisphere proportions were then averaged for each mouse.

We subtracted the matrices of estimated densities to visualize the difference of two-dimensional kernel density estimations and re-plotted these values as a heat map using the geom_tile function of ggplot2 (version 3.4.4; RRID:SCR_019186). For plotting of neuron locations on the rostrocaudal axis, we used the jitter function from base R (version 4.3.2, RRID:SCR_001905) using an amount of 0.075 mm to eliminate overplotting between the slices spaced 0.150 mm apart. To encircle the hindlimb field in this figure, we used the two-dimensional kernel density estimation function kde2d from the MASS package (version 7.3-60.0.1, DOI: 10.32614/CRAN.package.MASS) to draw a contour encompassing 99% of all labeled nuclei. Unpaired two-sided t-tests were performed for most quantities on each biological replicate (mouse). The data were not normal for counts of FG(+) neurons, so we used two-sided Wilcoxon signed-rank tests for each biological replicate (mouse), paired if making a within-mouse comparison, and unpaired if not. In both cases, no significant sex differences were observed.

## IV: Quantification and Statistical Analysis

### Anatomical Quantities

Sample sizes were determined using the variance and effect sizes from existing literature, and *a priori* sample size determination was not used. In subsequent statistical tests, n is defined as the number of biological replicates, which can be derived from individual mice or replicates with pooled mice. All statistical tests were performed using base R^133^ (version 4.3.2, RRID:SCR_001905) in RStudio^134^ (Build 494; RRID:SCR_000432). Details of individual statistical tests sufficient for reproduction are included in both the main text and figure legends. Exclusion criteria are defined in each relevant section of the methods. While most experiments were paired within-mouse, if relevant, group assignments were random with equal proportions by sex and cage. We did not detect sex differences within our analysis, but the experiments were not powered to detect such differences to maintain affordable project costs. This is an important limitation of the data presented in this study.

For within-mouse comparisons, paired two-tailed t-tests were performed. If examining proportions, one-sample t-tests were performed with μ = 0.5, representing a test against a null hypothesis of equal proportions. Anatomical quantities were assumed to be normally distributed, but in most cases, low sample sizes prevented formal testing of this assumption. Where relevant, we applied Bonferroni corrections for multiple comparisons. In figures, center was reported as mean or median, and if mean, precision was indicated visually by standard error of measurement. Dispersion was quantified using standard deviation.

The data in experiments examining neural regeneration were not normally distributed, as in existing studies of cervical innervation following hemisection.^103,104,106^ For these comparisons, we used two-sided Wilcoxon signed-rank tests. Dispersion was visualized with box and whisker plots, and precision was not indicated in figures because the statistical framework was based on rank comparison.

### Gene Expression Statistics

False discovery rates for differential expression analyses were determined using the default DESeq2 pipeline^160^ (version 1.42.0; RRID:SCR_015687). P-values for rank-rank hypergeometric overlap were computed using the default model in the RRHO package^97^ (version 1.42.0; RRID:SCR_014024). P-values for GSEA were determined using clusterProfiler^164^ (version 4.10.0; RRID:SCR_016884) package with a p-value cutoff of 0.01 and Benjamini-Hochberg correction for multiple comparisons.

## Acknowledgements

We are grateful to Rickard Sandberg, Christoph Ziegenhain, Etsuo Susaki, Katherine Stewart, Hanseul Yang, Jeffrey Macklis, Florence Bareyre, Arko Ghosh, Vibhu Sahni, Julia Kaiser, Natalie Russi, Edmund Hollis, Kelly Yee, Jonathan Esty, Irene Duba, Kip Lacey, Abdel El Manira, Christina Pressl, Kasia Turbek, Ivarine Rose, Alan Umfress, Rune Berg, Claudia Kathe, Christopher Crowley, Mor Alkaslasi, Claire Le Pichon, Ian Duguid, Chongyuan Luo, Salif Komi, Michael Riad, Kun Li, Vicky Moya, Hao Li, Francisca Martínez Traub, César Vargas, Matthew Davenport, Ivy Kosater, Meghana Rao, Elitsa Stoyanova, Bertrand Ottino-Loffler, Jeremy Nathans, Yuling Lin, Yutaka Yoshida, Jie Xing, Cuidong Wang, and Miro Koulnis for technical and administrative assistance and helpful discussions of this project and/or manuscript. Additionally, we wish to thank Svetlana Mazel of the Rockefeller University Flow Cytometry Resource Center, Connie Zhao of the Rockefeller University Genomics Resource Center, Ji-Dung Luo and Thomas Carroll of the Rockefeller Bioinformatics Resource Center, Tom Wiley, Stephanie Phillips, Leslie Diaz, Chaya Goodman, Jocelyn Guallpa, and David Powell of the Rockefeller Comparative Bioscience Center, Jim Petrillo of Rockefeller’s Gruss Lipper Precision Instrumentation Technologies facility, and Pantelis Tsoulfas and Yania Martinez of The Miami Project to Cure Paralysis. Finally, we wish to thank Cori Bargmann, Priya Rajasethupathy, and Jack Martin for their valuable insight and for serving on B.F.’s thesis committee.

For this work, B.F. was supported by the National Institutes of Health (NINDS 1F31NS129336). E.F.S. was supported by the National Institutes of Health (NINDS R01NS129902, R21NS133927). Spinal cord atlas generation was supported by the National Institutes of Health grant NIMH U19MH114830 to H.Z. N.H. was supported by the Howard Hughes Medical Institute (HHMI). O.S. was supported by the National Institutes of Health (NINDS R01NS047718, R01NS073857).

## Author contributions

B.W.F. and E.F.S conceived of the study and designed the experiments, with E.F.S. supervising and both providing funding. B.W.F. performed all investigation, formal analysis, data curation, and visualization unless otherwise noted below. I.T. wrote software and provided data curation for the analysis of the BDA tracing experiment. O.S. performed and funded BDA tracing experiments and provided resources for imaging. T.C.M. aided in methodological development for CUBIC tissue clearing and wrote software for the computational environment and image storage hardware to facilitate lightsheet imaging. M.H-J. aided in methodological development for Smart-Seq3 experiments. Y.G. performed the spinal cord data integration, with B.T., Z.Y., and H.Z. (AIBS) providing supervision and resources to this end. H.Z. (AIBS) and Z.Y. additionally provided funding for these experiments.

S.W.W. provided project administration for these efforts. H.Z. (CSHL) and C.F. performed MAPseq reverse transcription and sequencing and aided in methodological design, with H.Z. supervising, performing data curation, and providing resources to this end. B.C. performed vTRAP pulldowns and E.F.S. performed vTRAP dissections, with both providing methodological support. N.H. provided resources, funding, and supervision of the experiments and aided in conceptualization.

B.W.F. wrote the original draft. All authors reviewed the manuscript, with B.C., T.C.M, M.H-J., H.Z. (CSHL), Y.G., S.W.W, B.T., O.S., N.H., and E.F.S. contributing feedback that substantially altered the manuscript in its current form.

---

**Fig S1:**
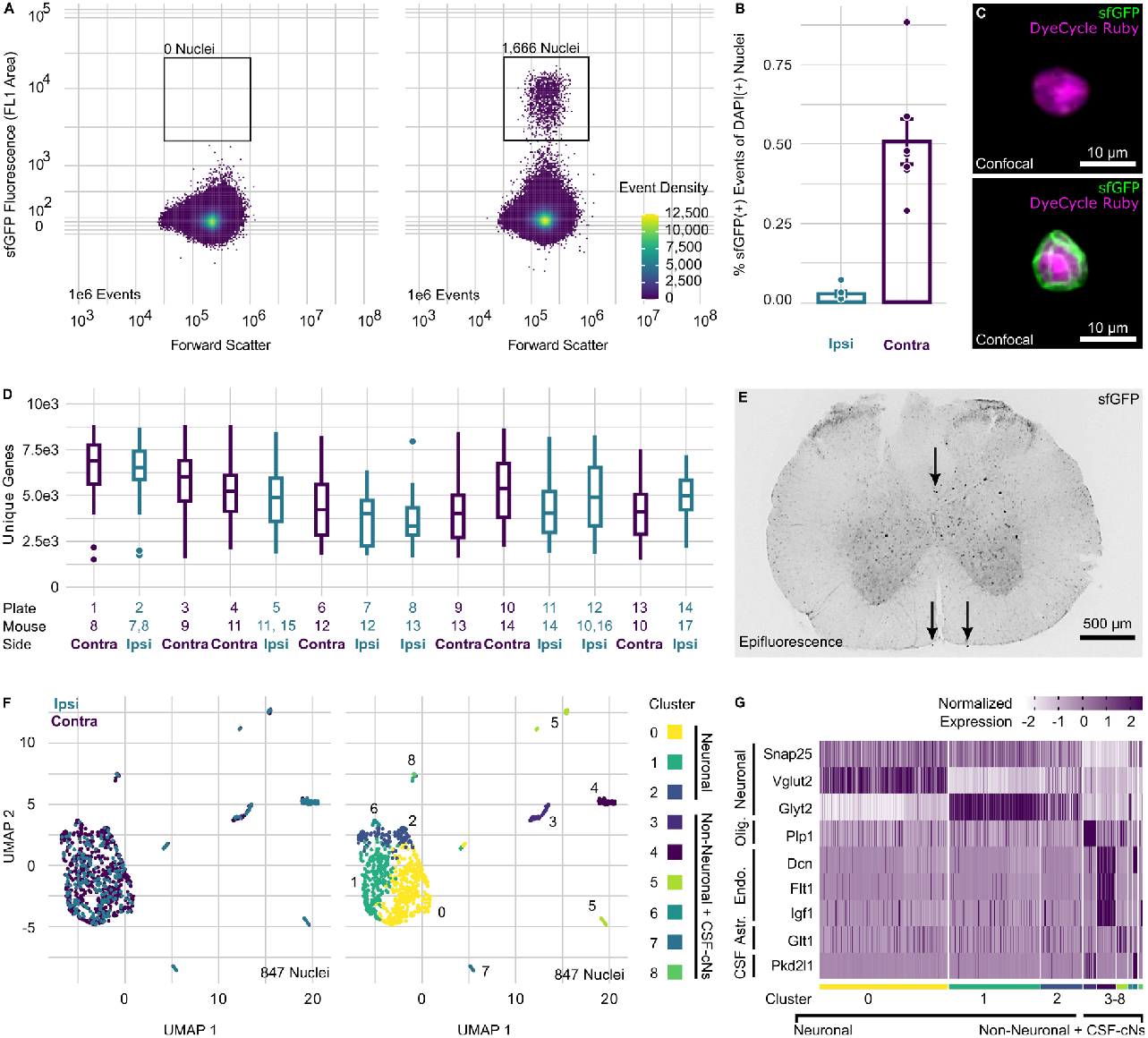
Supporting information for Smart-seq3 sequencing of neurons post-synaptic to the CST (related to Fig 2) **(A)** Example FACS density plots of single nuclei from the whole cervical spinal cord used for sfGFP gating. A negative control uninjected littermate is plotted on the left and an injected littermate on the right. The y-axis signifying the native nuclear sfGFP signal is scaled biexponentially. **(B)** A bar plot showing the percentage of sfGFP(+)-gated nuclei, with the percentage sfGFP(+) for the ipsilateral side samples in teal and the contralateral in purple. **(C)** Images of individual sfGFP(-) and sfGFP(+) nuclei form the sort in (A). **(D)** Box and whisker plots of unique genes detected in neuronal nuclei from each plate, with ipsilateral plates in teal and contralateral in purple. Outliers are shown as individual dots. **(E)** An example image of a cervical spinal cord slice with native nuclear sfGFP fluorescence. Arrows denote oligodendrocyte nuclei in the dorsal CST above and two meningeal nuclei on the bottom. The brightness of the representative image was adjusted in each channel so that tissue was visible via autofluorescence. **(F)** UMAPs of sorted sfGFP(+) nuclei. In the left panel, nuclei are colored by the side of origin, with ipsilateral in teal and contralateral in purple. On the right, the nuclei are clustered into eight transcriptomic groups and color-coded according to their group. **(G)** A heatmap showing normalized neuronal and non-neuronal marker gene expression. Each column represents a single nucleus. Nuclei are grouped by clusters.

**Fig S2:**
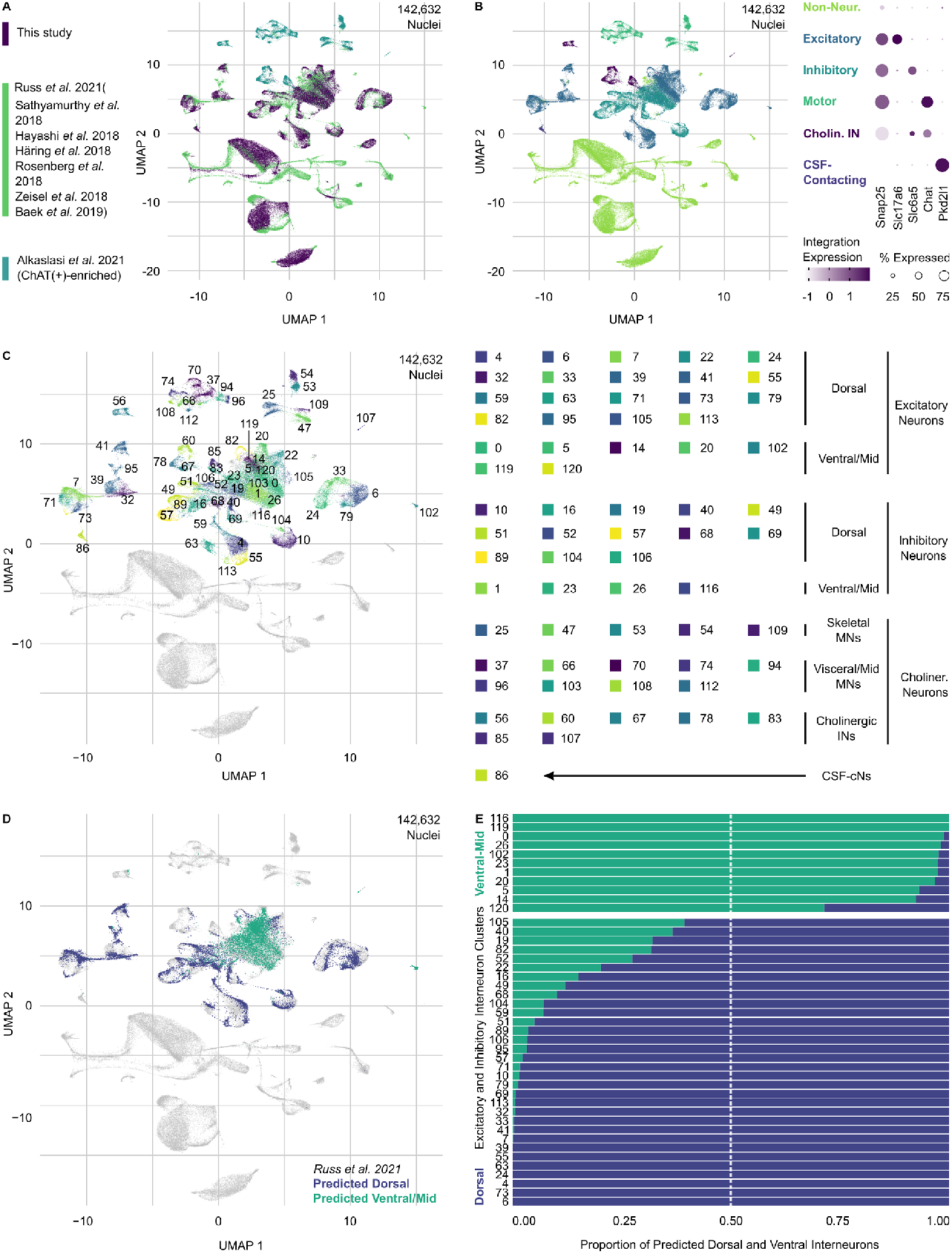
Supporting information for integrated atlas of spinal cord cell types (related to Fig 2) **(A)** A UMAP of 142,632 integrated spinal cord single-cell and single-nucleus transcriptomes colored according to their study of origin. **(B)** A UMAP of coarse-grained clusters in the entire data set. **(C)** A UMAP of neuronal transcriptomic clusters labeled and colored according to cluster membership. Non-neuronal nuclei in the data set are colored gray. **(D)** A UMAP showing the location of nuclei from Russ *et al*. 2021 colored according to the paper’s designation of the nuclei as located dorsally or middle-to-ventrally. Other nuclei in the data set are colored gray. **(E)** A bar plot showing the fraction of nuclei locations from Russ *et al*. 2021 in each cluster and each cluster’s putative location.

**Fig S3:**
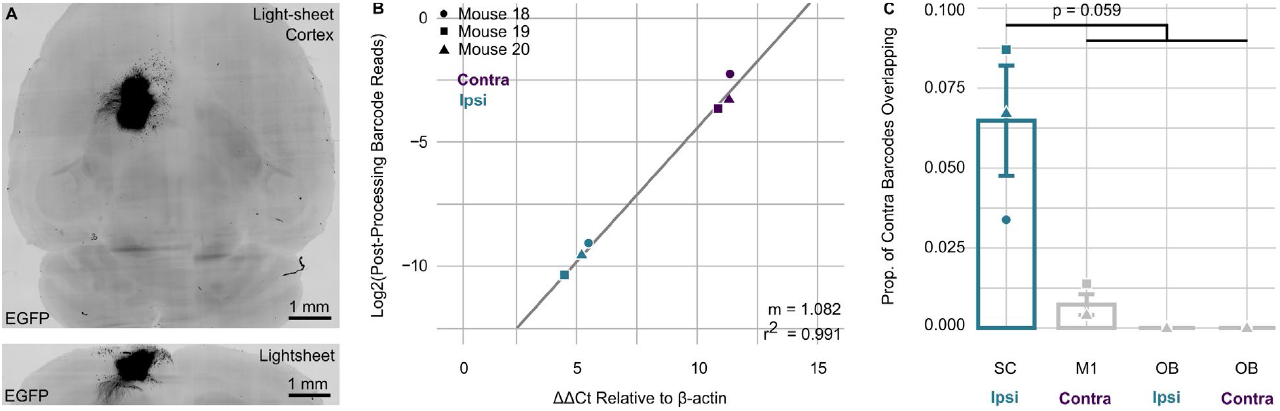
MAPseq Supporting Information (related to Fig 4) **(A)** MAPseq Sindbis virus injection sites, which express EGFP. On the top is a maximum-intensity z-projection on the horizontal plane with native EGFP fluorescence. On the bottom, a 100 μm maximum-intensity z-projection on the coronal plane. The brightness of representative images was adjusted so that tissue was visible via autofluorescence. **(B)** A scatter plot of Log2-transformed quantifications of MAPseq signal strength among spinal cord samples. On the x-axis are ΔΔCt values relative to β-actin from each sample’s initial quantification of *EGFP* transcripts, which is co-expressed with barcode transcript. On the y-axis are log2-transformed barcode sequence counts after quality control. **(C)** A bar plot showing the proportion of contralateral spinal cord barcodes also detected in other regions. In teal, the proportion of contralateral barcodes detected on both ipsilateral and contralateral sides of the spinal cord. In gray, the proportion of contralateral barcodes detected in the areas that ought not to have barcodes according to prior tracing studies (p = 0.059, n = 3, one-sample two-sided t-test, μ = 0.5).

**Figure S4:**
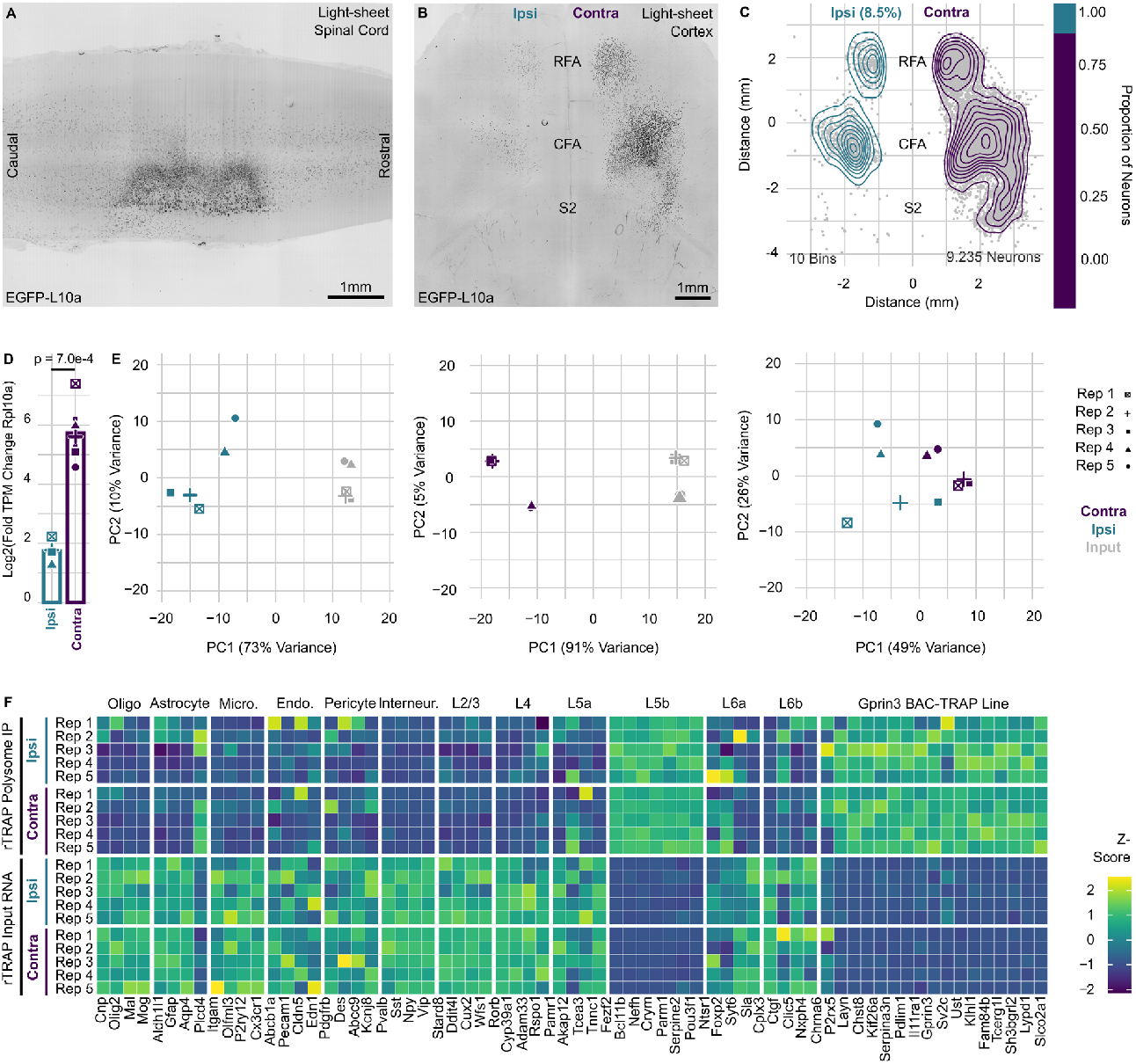
Isolation of translating mRNAs in ipsilateral and contralateral CSNs (related to Fig 6) **(A)** A maximum-intensity z-projection of the cervical enlargement and unilateral injection site EGFP-L10a expression on a coronal plane. **(B)** A maximum-intensity z-projection of the motor cortex on a horizontal plane, with corticospinal neurons expressing native EGFP-L10a fluorescence. The brightness of representative images was adjusted so that tissue was visible via autofluorescence. **(C)** A scatter plot of EGFP-L10a(+) corticospinal neurons on a horizontal plane. Density plots (10 bins) of neuronal density are overlaid, with ipsilateral in teal and contralateral in purple. A stacked bar plot to the right indicates the proportions of annotated ipsilateral and contralateral neurons in that sample. **(D)** Bar plots of Log2(fold-change TPM) over unprecipitated input RNA of *Rpl10a* transcripts (p = 7.0e-4, n = 5, paired two-sided t-test), with ipsilateral TPMs in teal and contralateral in purple. **(E)** Scatter plots of the first two principal components of each replicate. In the left-most panel, libraries from the ipsilateral projection-enriched immunoprecipitated polysomes in teal are compared to their matched unprecipitated input RNA samples in gray. Next, the same comparison for the contralateral projection-enriched libraries in purple relative to their unprecipitated input samples in gray. The right-most plot compares the ipsilateral immunoprecipitated samples in teal and the contralateral ones in purple. **(F)** A heatmap of cortical cell type marker gene z-scores for all samples in the experiment. Z-scores were calculated among all samples of a given gene. The first five blocks of columns are canonical marker genes for non-neuronal cell types of the cortex. The subsequent seven blocks are canonical marker genes for each classical cell type of the motor cortex, including Layer 5b. Finally, the last block of rows represents genes from a *Gprin3* BAC TRAP line, which labels Layer 5b corticospinal neurons.

**Figure S5:**
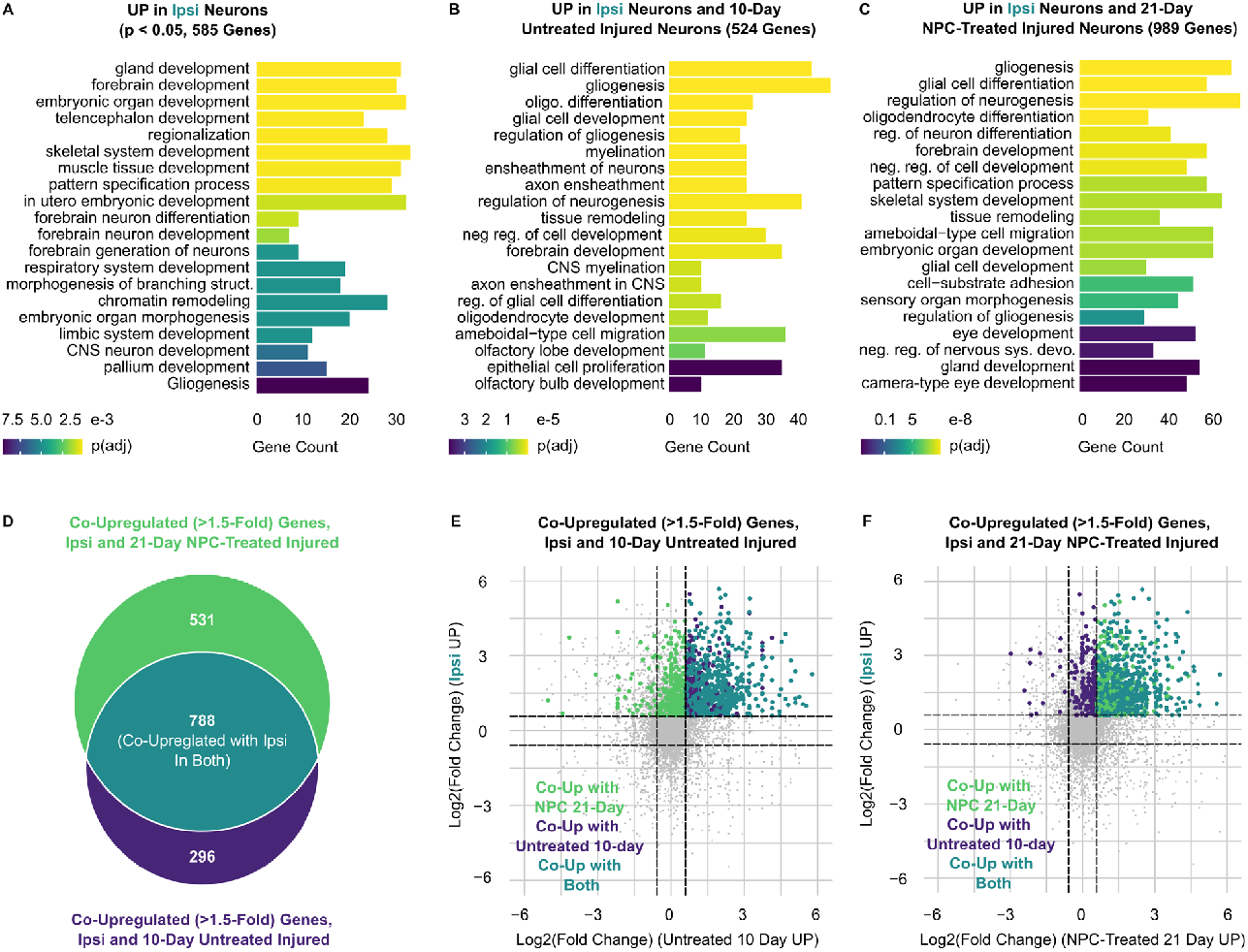
Gene Ontology (GO) analysis of differentially expressed gene sets (related to Fig 5) **(A)** Biological process terms that are significantly over-represented in the IP-CSN-enriched genes with an FDR < 0.05. **(B)** Biological process terms that are significantly over-represented in both the IP-CSNs and the untreated corticospinal neurons ten days after injury. **(C)** Biological process terms that are significantly overrepresented in both the IP-CSNs and NPC-treated corticospinal neurons 21 days after injury. **(D)** A Venn diagram of the genes used for enrichment analysis in (B) and (C) compares whether the upregulation shared by the IP-CSNs and the injured neurons is similar in NPC-treated mice at the 21-day time point and the untreated mice at the 10-day time point. **(E)** A scatter plot of Log2(fold) changes compares molecular changes between the ipsilateral and contralateral CSN populations and those in injured but untreated mice ten days post-injury. Genes are color-coded according to the Venn diagram in (D) to examine whether the upregulation shared by the IP-CSNs and the injured neurons is similar in NPC-treated mice at the 21-day time point and the untreated mice at the 10-day time point. **(F)** A scatter plot of Log2(fold) changes comparing molecular changes between the ipsilateral and contralateral CSN populations and those in injured and NPC-treated mice at 21 days post-injury. Genes are color-coded according to the Venn diagram in (D) to examine whether the upregulation shared by the IP-CSNs and the injured neurons is similar in NPC-treated mice at the 21-day time point and the untreated mice at the 10-day time point.

**Figure S6:**
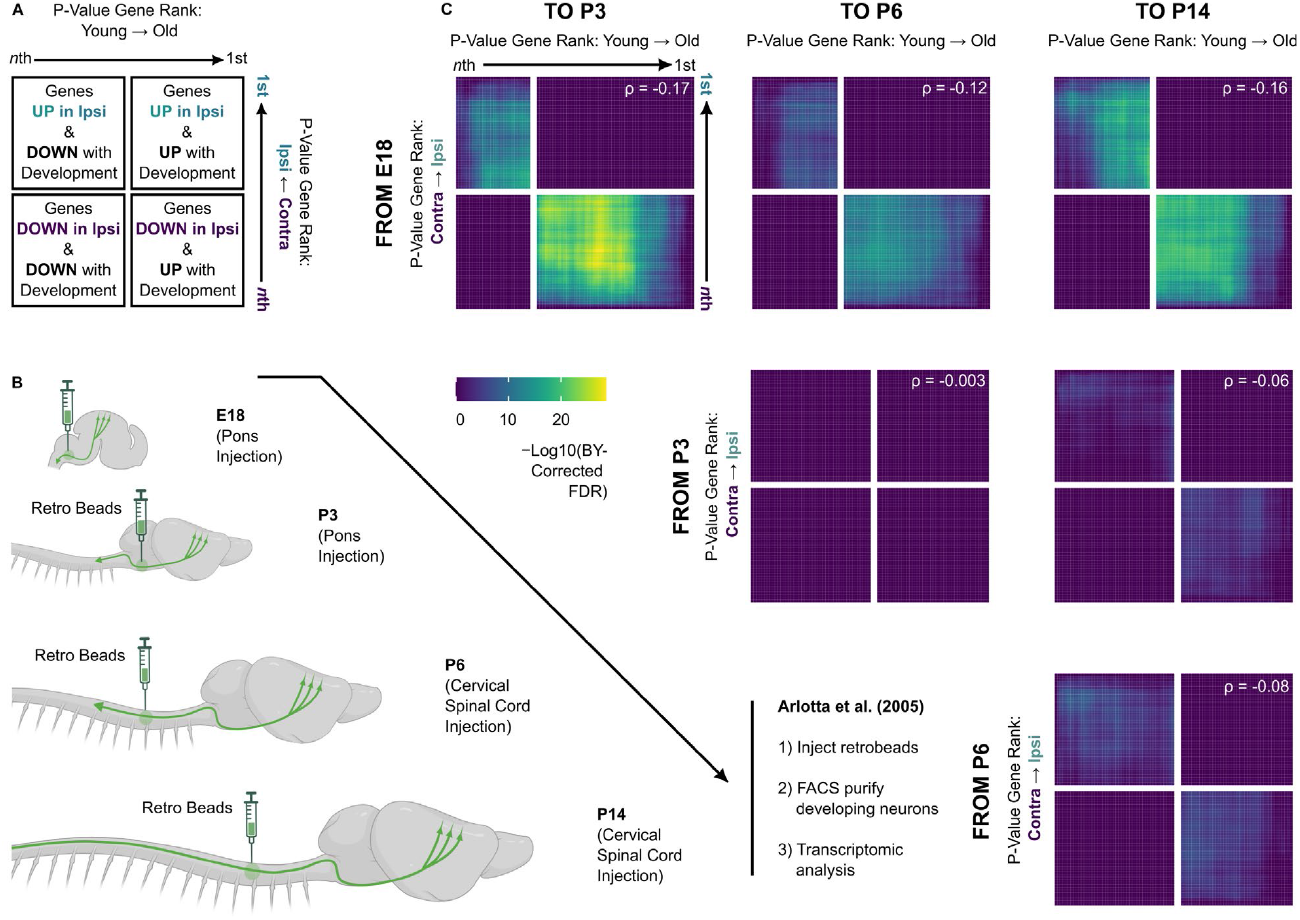
IP-CSNs bear an embryonic-like molecular state (related to Fig 6) **(A)** A schematic of Rank-Rank Hypergeometric Overlap (RRHO) analysis as performed in subsequent panels. **(B)** A schematic of Arlotta *et al*. 2005’s experiments examining the transcriptomes of corticospinal neurons throughout development. **(C)** RRHO analysis comparing the differential expression of ipsilateral vs. contralateral CSN populations with those of Arlotta *et al*. 2005. The figure is arranged so that each panel represents differential expression from one time point in development to each later time point. For example, the top row of correlations compares our vTRAP differential expression data with the molecular changes occurring from E18 to P3 in the left-most panel, E18 to P6 in the middle panel, and E18 to P14 in the right-most. Negative Spearman’s ρ values indicate differential expression discordant with that of ipsilateral projection-enriched neuronal populations as the neurons develop. FDR values are expressed using the Benjamini-Yekutieli method for multiple comparison correction.

**Fig S7:**
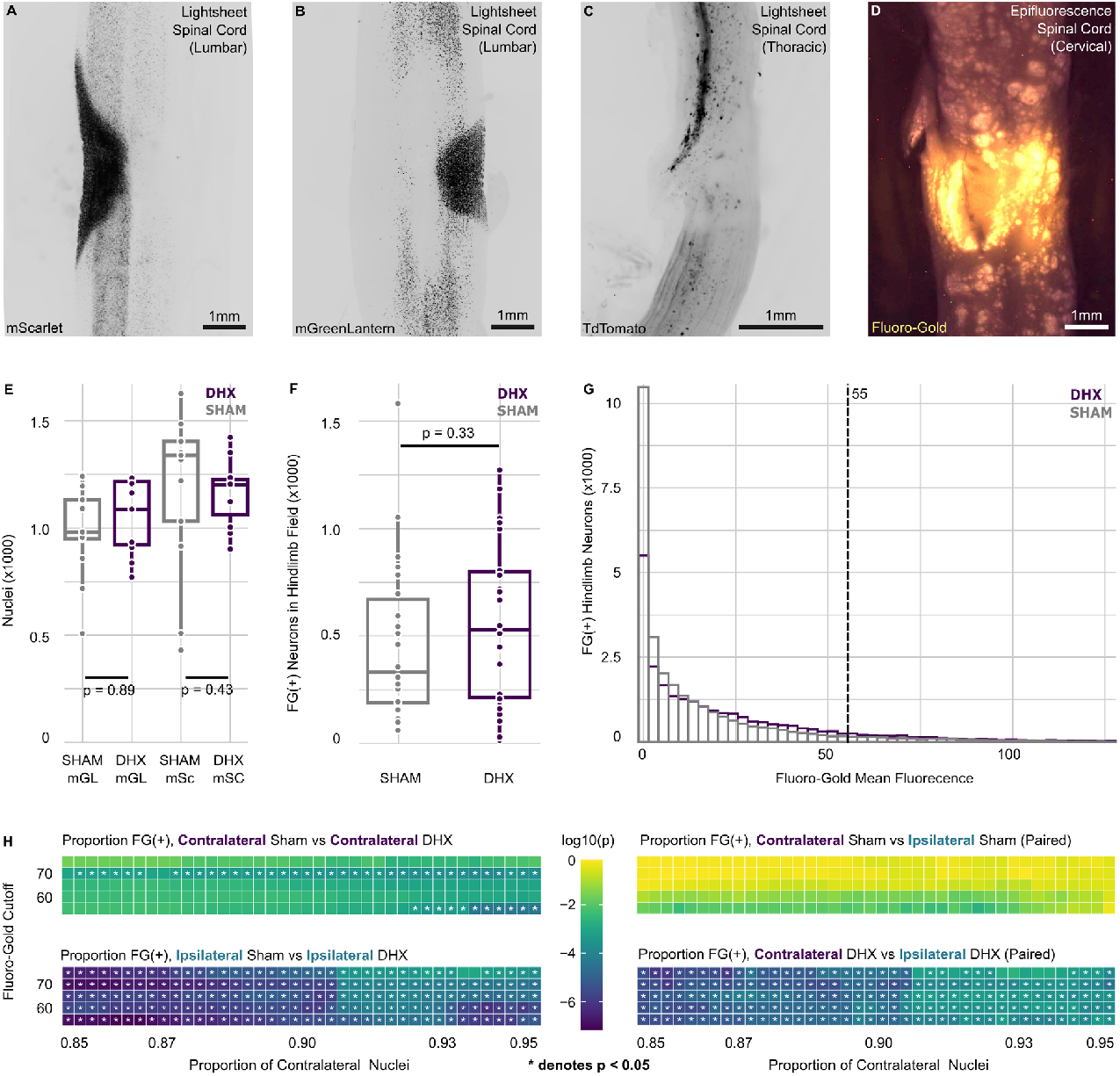
Supporting information for triple tracing of corticospinal neurons after dorsal hemisection (related to Fig 7) **(A)** A maximum-intensity z-projection on a coronal plane of a cleared mScarlet-H2B-retro injection site expressing native mScarlet. **(B)** A maximum-intensity z-projection on a coronal plane of a cleared mGreenLantern-H2B-retro injection site expressing native mGreenLantern. **(C)** A maximum intensity z-projection on a sagittal plane of a cleared dorsal hemisection. AAV1-Cre was injected into the hindlimb cortex of an Ai14 tdT reporter mouse such that the severed CST was visible as native tdT signal **(D)** An image of a cleared cervical spinal cord fluorogold injection site from a coronal perspective. UV light did not penetrate the cleared tissue in a light-sheet microscope, so this image of native FG fluorescence was taken on an epifluorescence microscope. The brightness of representative images was adjusted in each channel so that tissue was visible via autofluorescence. **(E)** Box plots showing total fluorescent nuclei counts among all mice in both sham and hemisected groups from both fluorophores (p = 0.89 mGL and 0.43 mSc, n = 11 DHX, 13 sham, unpaired two-sided t-test). Hemisected mice are displayed in purple and sham in gray. **(F)** Box plots showing total FG(+) neuron counts, irrespective of whether they coincided with lumbar projecting nuclei (p = 0.33, n = 11 DHX, 13 sham, unpaired two-sided t-test). **(G)** A histogram shows the FG signal distribution among all retrogradely traced lumbar projecting nuclei in sham and hemisected mice. The dotted line at 55 represents the cutoff used in Figure 7. **(H)** Heat maps showing the analysis results in Figure 7C for various combinations of the two threshold values. In each, the rows represent FG cutoffs from 55–75. Columns represent ipsilateral fluorophore cutoffs such that 85–95% of nuclei are designated contralateral. The two left heat maps are the results of Wilcoxon signed-rank tests of the share of FG(+) neurons in hemisected mice vs sham, in contralateral neurons (top) and ipsilateral neurons (bottom). The two right heat maps are paired Wilcoxon signed-rank test between ipsilateral and contralateral neurons in the sham mice (top) and in the hemisected mice (bottom). Asterisks denote p-values less than 0.05.

